# A feedback loop of conditionally stable circuits drives the cell cycle from checkpoint to checkpoint

**DOI:** 10.1101/654863

**Authors:** Dávid Deritei, Jordan Rozum, Erzsébet Ravasz Regan, Réka Albert

## Abstract

We perform logic-based network analysis on a model of the mammalian cell cycle. This model is composed of a Restriction Switch driving cell cycle commitment and a Phase Switch driving mitotic entry and exit. By generalizing the concept of stable motif, i.e., a self-sustaining positive feedback loop that maintains an associated state, we introduce the concept of conditionally stable motif, the stability of which is contingent on external conditions. We show that the stable motifs of the Phase Switch are contingent on the state of three nodes through which it receives input from the rest of the network. Biologically, these conditions correspond to cell cycle checkpoints. Holding these nodes locked (akin to a checkpoint-free cell) transforms the Phase Switch into an autonomous oscillator that robustly toggles through the cell cycle phases G1, G2 and mitosis. The conditionally stable motifs of the Phase Switch Oscillator are organized into an ordered sequence, such that they serially stabilize each other but also cause their own destabilization. Along the way they channel the dynamics of the module onto a narrow path in state space, lending robustness to the oscillation. Self-destabilizing conditionally stable motifs suggest a general negative feedback mechanism leading to sustained oscillations.

## Introduction

Bottom-up mechanistic models of biological regulation are becoming increasingly adept at describing and predicting complex cellular behavior^1–8^. As qualitative experimental data on these regulatory interactions are more abundant than kinetic information, modeling frameworks that leverage qualitative data are often used to probe the behavior of large regulatory networks. Boolean models, in particular, assume that the essential logic by which regulatory interactions drive cell behavior is adequately described by a network of molecular components (e.g. mRNA, transcription factors, signaling proteins), where the activity of each is described by a binary variable. Boolean models have been shown to faithfully reproduce cellular phenotypes and behaviors in cells from all domains of life^9–19^. Overall, the successes of Boolean modeling reveal that complex, context-dependent cellular behaviors generally do not rely on narrowly tuned kinetic parameters. Rather, the network topology and combinatorial logic of regulatory interactions are key determinants of cellular function.

The idea that the structure of a regulatory circuit is key to its dynamic behavior inspired a series of methods to link features of network topology to biological insight^20–30^. For Boolean models, the focus is on identifying stable states or sustained oscillatory behaviors, called *attractors*; these correspond to cellular phenotypes. Thus, identifying all the attractors of a Boolean model is a critical step in probing the model’s ability to reproduce the phenotypic repertoire and responsiveness of cells. A recently introduced method to connect the regulatory network to the attractor repertoire of biological systems encodes the regulatory logic in an *expanded network*^*31*^ by the inclusion of a separate *on* and *off* state for each node, as well as composite nodes to represent input combinations that must work together (i.e., AND gates). Analysis of the expanded network can identify minimal building blocks that dictate the system’s dynamics. Subgraphs of the expanded network termed *stable motifs* indicate node values that stabilize independently of the rest of the network; stable motifs consequently trap the system’s dynamics into specific regions (or sub-spaces) of the state space. The sequential lock-in of stable motifs restricts the system’s dynamics until it reaches an attractor. A counterpart to stable motifs, *oscillating motifs*, identify node subsets that cannot achieve a stable state, and thus give rise to oscillating or complex attractors^31,32^.

An advantage of logic-based structural analysis is that it can reveal the dynamical building blocks, or *modules*, that cellular regulation uses to drive distinct outcomes. Indeed, multiple lines of evidence suggest that cell-wide interaction networks are made of small tightly connected modules, linked into a hierarchy of looser modules at each scale^33–37^. Cells respond to their environment with discrete combinations of specific functions and often break down one functional module at a time^38,39^. Deritei et al argued in 2016 that cellular regulatory networks exhibit *dynamical modularity*^*17*^: they are composed of a hierarchy of coupled multi-stable modules, each of which is responsible for a discrete set of mutually exclusive phenotypic outcomes, such as survival vs. apoptosis (programmed cell death), cell cycle progression vs. cell cycle arrest. The global fate of cells then consists of distinct combinations of states created by isolated modules. Dynamical modularity also explains biological rhythms generated by coupled multi-stable modules. Indeed, Deritei et al. construct a mammalian cell cycle model built from two multi-stable modules to showcase the dynamical modularity of the cell cycle. The two modular switches -- the *Restriction Switch* responsible for cell cycle commitment and the *Phase Switch* in control of entry and exit from mitosis (cell division) -- toggle each other such that the cell state follows a repeating sequence of distinct combinations of the switches’ stable states. Moreover, the attractors of a significantly larger five-module Boolean model of cell cycle coordination with growth factor signaling and show similarly modular dynamics^19^.

We performed logic-based structural analysis of the cell cycle model of Deritei et al. to identify the stable motifs that drive the attractors of each individual module and pinpoint the oscillating motif responsible for the cell cycle. A possible mechanism for a dynamically modular cycle is the existence of long, cross-module negative feedback loops in which a stable state of one switch toggles the state of a second switch, which then destabilizes the first. While inter-module feedback is indeed present, here we show that the Phase Switch *alone* has internal structural features capable of generating a robust cell cycle oscillation.

## Background and key concepts

Boolean models of biological regulatory systems have been described in several recent review articles^40–43^. Here we review key concepts of Boolean modeling. Boolean models assume that the activity of each element can be described by two qualitative states: 1 (interpreted as on, or active) and 0 (interpreted as off, or inactive). The future state of each node depends on the current state of its regulators (parents in the network) in a way described by a *regulatory function x*\* = *f*_*x*_ (*Par(x)*), where for simplicity the state of the node is denoted with the node name, *x*\* is the next state of node *x, f*_*x*_ is the regulatory function of *x* and *Par(x)* represents the parent nodes of (nodes with edges pointing toward) *x*. The order in which the nodes of a Boolean model are updated can significantly impact the emergent dynamical trajectories. In this paper we focus on two update schemes: synchronous update (used in the Deritei et al. model^17^ of the cell cycle) and general asynchronous update. In *synchronous update* all nodes are updated at the same time and their next state is determined by the previous state of the system. Synchronous update yields deterministic trajectories. In the *general asynchronous update* scheme, the next node to be updated is chosen randomly, which adds a large degree of stochasticity into the emergent trajectory of the model.

All trajectories of Boolean systems ultimately lead into attractors. An attractor is a single state, or a set of states, that the system cannot leave. Attractors that are single states are called fixed-point attractors (or steady states). Fixed points of Boolean models of biological systems represent stable phenotypes. Attractors that contain multiple states are called complex attractors (or limit cycles in case of synchronous update). Limit cycles usually correspond to oscillations such as the cell cycle. The *state transition graph (STG)* is the network representing all the possible transitions between the states of a system. Fixed point attractors correspond to sink nodes (i.e. nodes lacking outgoing edges) of the STG. Complex attractors correspond to terminal strongly connected components of the STG (i.e. there are no state transitions that lead out of the attractor). The fixed points of Boolean models are independent of update scheme, but the complex attractors may change or disappear when using different types of update.

Much of our analyses rely on the construction and properties of an *expanded network*^*9,31,32,44,45*^. The expanded network of a Boolean system encodes the causal relationships between node states reflected in the regulatory functions. It consists of two “virtual nodes” for each node (one for each of the two possible states) and “composite nodes” that embody AND gates among two or more node states. An edge from a virtual node to a composite node indicates that the virtual node is a necessary condition for states described by the composite node. An edge from any node to a virtual node indicates that the parent node is a sufficient condition for the state represented by the child node. Figure 1 indicates the expanded network that corresponds to the network and regulatory functions of a hypothetical example.

**Figure 1.**
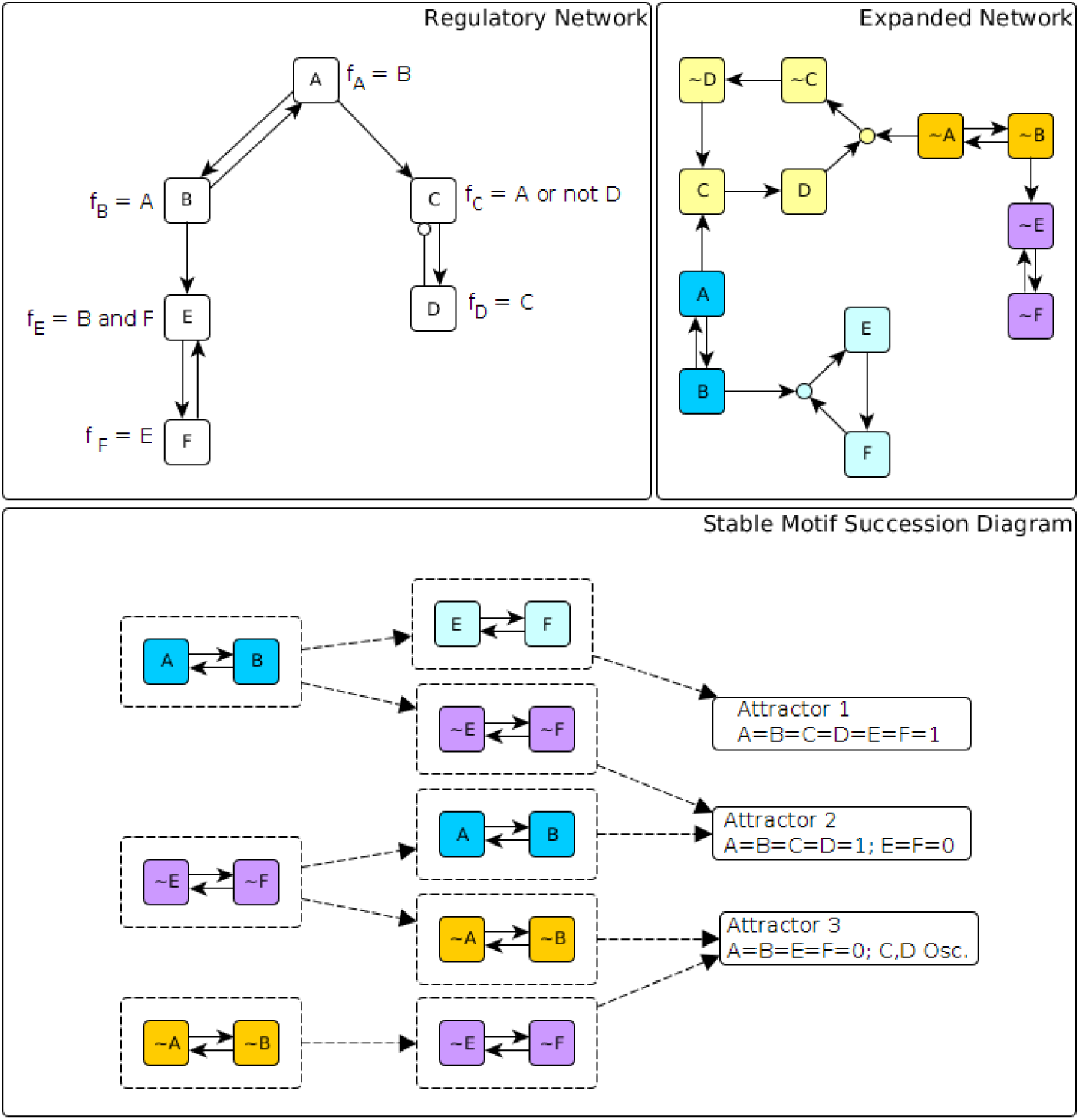
Illustration of the expanded network, stable motifs and conditionally stable motifs of a hypothetical network. In the network (shown on the top left panel) edges with terminal arrows indicate positive regulation and edges that end in open circles indicate negative regulation. The regulatory function of each node is indicated next to the node. The expanded network (top right panel) includes two virtual nodes for each node: the virtual node marked by the node name indicates the on (1) state of the node while the virtual node marked by the node name preceded by ∼ indicates the off (0) state of the node. The expanded network also includes composite nodes (small filled circles) for each AND gate, e.g. the composite node with light blue background indicates the AND gate in the regulatory function of node E. The positive feedback loop between A and B yields two stable motifs, one corresponding to the on state of both nodes (shown in blue) and one corresponding to the off state of both nodes (shown in orange). The positive feedback between E and F can sustain their off state (stable motif shown in purple) but the on state of E and F can only be sustained if B is on. Thus, the virtual nodes E and F form a conditionally stable motif conditioned on B (light blue). The negative feedback between C and D leads to sustained oscillation of these two nodes (indicated by the cycle in yellow) if A=0. If A=1 C will also converge to 1 (see the edge from A to C in the expanded network). The stable motif succession diagram shown in the bottom panel indicates the possible sequences of successive stabilization of the three stable motifs and of the conditionally stable motif as well as the resulting attractor repertoire of the system.

Within an expanded network, there are subgraphs that capture important dynamical features of the underlying system. Of particular interest here is the class of subgraphs called *stable motifs*^*31*^. A stable motif is a subgraph of an expanded network that satisfies four properties: 1) it is *strongly connected* (there is a path between every pair of nodes in the subgraph), 2) it is *consistent* (all represented conditions can be simultaneously satisfied), 3) it is *composite-closed* (if a composite node is in the subgraph, so too are all its virtual node parents), and 4) it is *minimal* (it contains no subgraphs, other than itself, satisfying the first three properties). A stable motif determines a (non-empty) region of the state space from which dynamical trajectories cannot escape. In general, this region constrains some of the system variables (including the variables of the stable motif), but not others. The confinement of the constrained variables will be independent of the unconstrained variables even when these unconstrained variables are externally manipulated.

Another important class of expanded network subgraph is the oscillating motif. Like stable motifs, oscillating motifs are strongly connected, composite-closed subgraphs. Unlike stable motifs, however, oscillating motifs violate the consistency criterion in that every virtual node in the subgraph has its negation in the subgraph as well. Furthermore, an oscillating motif may not contain any stable motifs as subgraphs. An oscillating motif is so-named because its presence implies the existence of a complex attractor in which all of the nodes represented in the motif oscillate^31^.

By substituting the node states of a stable motif into the Boolean regulatory functions and simplifying, one obtains reduced functions that describe the system evolution sufficiently long after the stable motif trap space has been entered. Using these rules, a new expanded network can be generated, in which additional stable motifs may be identified. The succession of stable motif identification and network reduction ends when there no longer are any stable motifs (when this occurs, the expanded network is either empty or it likely is an oscillating motif^31^). At each step in the succession, there may be several stable motifs; by considering all possible orders of stable motif selection, the attractors of the Boolean system can be enumerated. The bottom panel of Figure 1 illustrates this procedure and the resulting paths to the system’s three attractors.

In this work, we introduce a new generalization of stable motifs that allows node states to stabilize when a specified set of conditions are satisfied. In particular, we define “conditionally stable motifs” (CSMs) to be consistent, strongly connected components of the expanded network that are not composite closed. For a given CSM, we define the CSM conditions to be the set of virtual nodes that are external to the CSM but are parents of composite nodes internal to the CSM. A stable motif can be viewed as a CSM with an empty condition set. A CSM becomes a stable motif under the assumption that its conditions are satisfied. We provide a method to construct all CSMs in a Boolean network and implement this method on the Phase Switch Oscillator (see Methods). The number of CSMs can become very large, which complicates interpretation of the collective significance of a system’s CSMs. For this reason, we focus our attention on two classes of CSM: those with the fewest conditions and those that are as large as possible.

### Three stable motifs and four conditionally stable motifs determine the three fixed point attractors of the Phase Switch

The Deritei et al. cell cycle model^17^ contains two modules, the Restriction Switch (6 nodes), the Phase Switch (11 nodes) and three abstract nodes (Replication, Metaphase, 4N DNA), which represent regulatory modules of the cell that are not explicitly included the model. The Restriction Switch is bistable, with point attractors that correspond the before- and after states of the restriction point, and the Phase Switch is tri-stable, with point attractors that match the phases of the cell cycle. Specifically, the activity of the nodes of the Phase Switch in the three fixed points match those of cells in G0/G1, G2 phases of the cell cycle, and the Spindle Assembly Checkpoint (SAC), respectively (see Supplementary Table S1). As cells enter a division cycle, the Restriction Switch in control of this commitment toggles the Phase Switch by tipping it from its quiescent G0/G1 attractor into G2, a state that corresponds to the DNA damage (G2) checkpoint. This in turn resets the Restriction Switch to an uncommitted state. Passage of the G2 checkpoint tips the Phase Switch into mitosis, where it maintains a stable Spindle Assembly Checkpoint. Once the mitotic spindle is fully assembled -- an event that impacts the Phase Switch from the outside -- it resets to its initial state, the G0/G1 attractor. This, in turn, allows the Restriction switch to commit to another cycle.

In order to understand the structural causes of this modular toggle, we first focused on characterizing the Phase Switch. The interactions among the 11 nodes of the Phase Switch express either positive or negative regulatory effects (Figure 2). The Phase Switch is strongly connected; all nodes within the module can be reached from all other nodes via at least one directed path. It contains both positive and negative feedback loops of various lengths. Using the expanded network formalism to integrate the structure and regulatory functions of the Phase Switch (given in Supplementary Text S1) we identified three stable motifs, P0 to P2, as shown on Figure 2. Additionally, there are four conditionally stable motifs: conditionally stable motifs P3 and P4 depend on the prior establishment of CyclinA=0. Conditionally stable motif P5 is a stable motif only if P1 is already established. Conditionally stable motif P6 is a stable motif only if P1 and P5 are already established.

**Figure 2.**
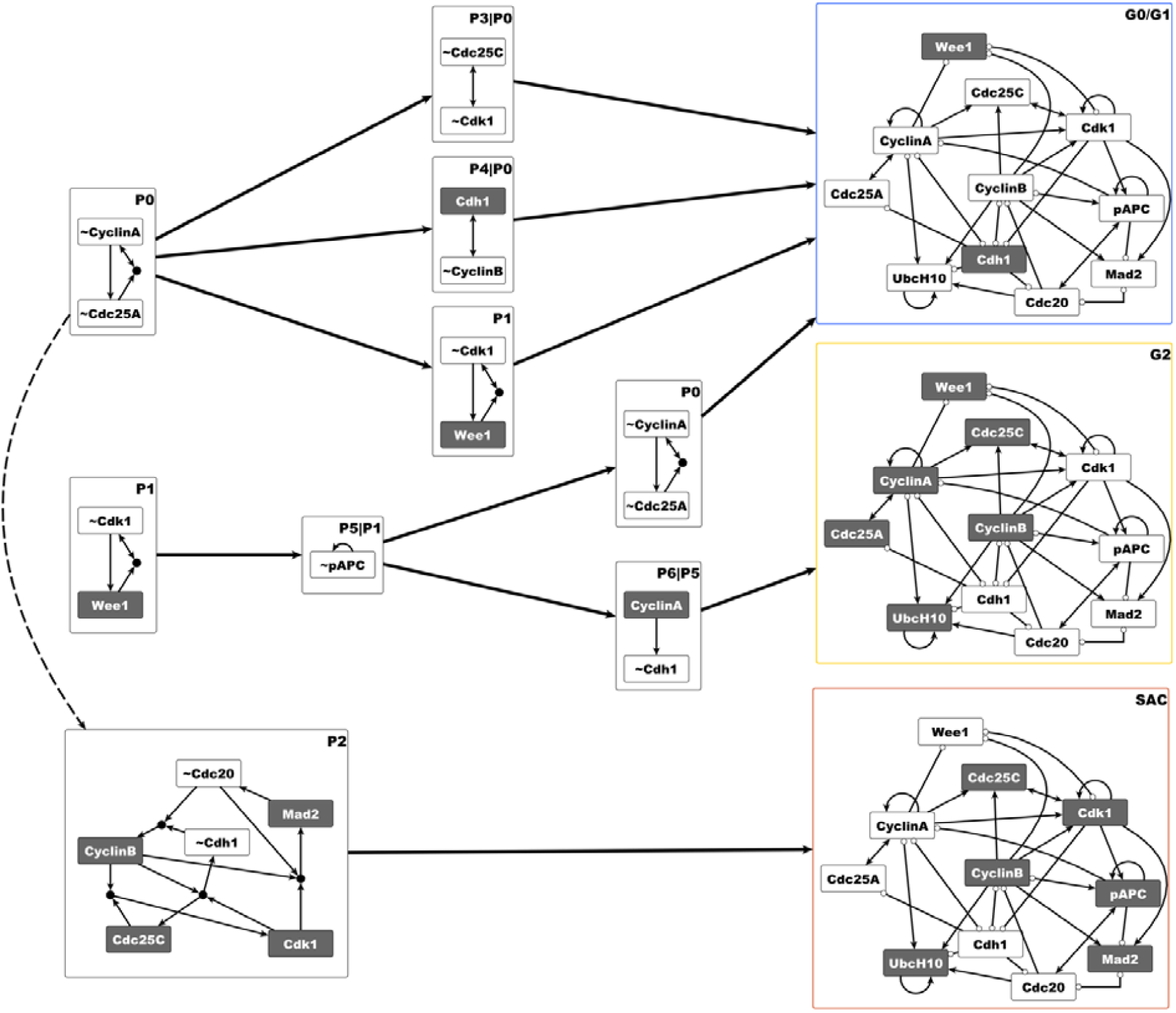
Distinct sequences of stable and conditionally stable motifs commit the Phase Switch to its attractor states. The stable motifs are shown in the expanded network formalism where a virtual node labeled by the name of the corresponding node represents the state 1 of the node; a virtual node labeled by the node name preceded by ∼ represents the state 0 of the node. The coloring of the virtual nodes conveys the same information: white background indicates that the node is off (0) and dark grey background indicates that the node is on (1). Stable motifs P0, P1 and P2 are the stable motifs of the Phase Switch. When substituting the states of P0 (i.e. CyclinA=Cdc25A=0) into the Boolean update functions of the Phase Switch, the resulting 6-node reduced system has four stable motifs (P1, P2, P3 and P4). Further reduction by substituting either P3 or P4 yields the G0/G1 attractor of the system (indicated in the network of the Phase Switch by the node colors, where dark grey means 1 and white means 0). The same attractor can also be found by successive substitution of P1, P5 and P0. The succession of P1, P5 and P6 yields the G2 attractor, and the P2 stable motif (with or without prior stabilization of P0, the former shown as a dashed arrow) yields the SAC attractor. The outline color of the three attractors corresponds to the color-code used for differentiating the three attractors in Deritei et al. 2016^17^.

The stable motif succession diagram of the Phase Switch (Figure 2) indicates that sequential stabilization of stable motifs and conditionally stable motifs underlies the three fixed point attractors previously identified by Deritei et al. The P1 and P2 stable motifs are mutually exclusive, as they contain opposite states of Cdk1. The P2 stable motif drives the SAC attractor (see Figure 2 and Supplementary Table S1). Each of the other two stable motifs is consistent with two attractors, depending on which other motifs stabilize. P0 is consistent with G0/G1 and SAC. P1 is consistent with G0/G1 and G2. The activation of each motif represents a decision point in the system trajectory. For instance, activation of the P0 motif prevents the system from entering the G2 state. Activation of the P1 motif represents a decision to not enter the SAC state, while activation of P2 always leads to the SAC state. Together, these three motifs are sufficient to determine the system’s ultimate steady state: G0/G1 is reached when P0 activates without activation of P2, G2 is reached when P1 activates without activation of P0, and SAC is reached when P2 activates.

### Stable motifs of the Phase Switch are conditionally stable motifs of the cell cycle model they are embedded in

In the full cell cycle model^17^ the nodes E2F1 and CyclinE of the Restriction Switch and the three abstract nodes regulate three nodes of the Phase Switch, namely Cdc25A, Wee1 and Mad2 (see Figure 3 and Supplementary Text S1). Because of these incident influences, the stable motifs of the Phase Switch are only conditionally stable in the context of the larger model. The stabilization of the P0 motif requires the OFF state of either E2F1 or CyclinE (see P0_CSM on Figure 3). The stabilization of the P1 motif requires the ON state of the Replication node, while the stabilization of P2 requires the simultaneous OFF state of Metaphase and ON state of 4N DNA. All of these nodes are in their required states for only specific phases of the cell cycle; for example E2F1 and CyclinE are off in the uncommitted state of the Restriction Switch. Because of this dependence on external regulators that are only transiently in the state necessary for stabilization, the P0, P1, P2 motifs of the Phase Switch cannot stabilize permanently. In a dividing cell, the period during which one of these motifs maintains its stability corresponds to a cell cycle checkpoint: the restriction point for P0, the G2 DNA damage checkpoint for P1, and the spindle assembly checkpoint (SAC) for P2^46^. Passage of each checkpoint changes the conditions around the Phase Switch such that the corresponding motif becomes unstable.

**Figure 3.**
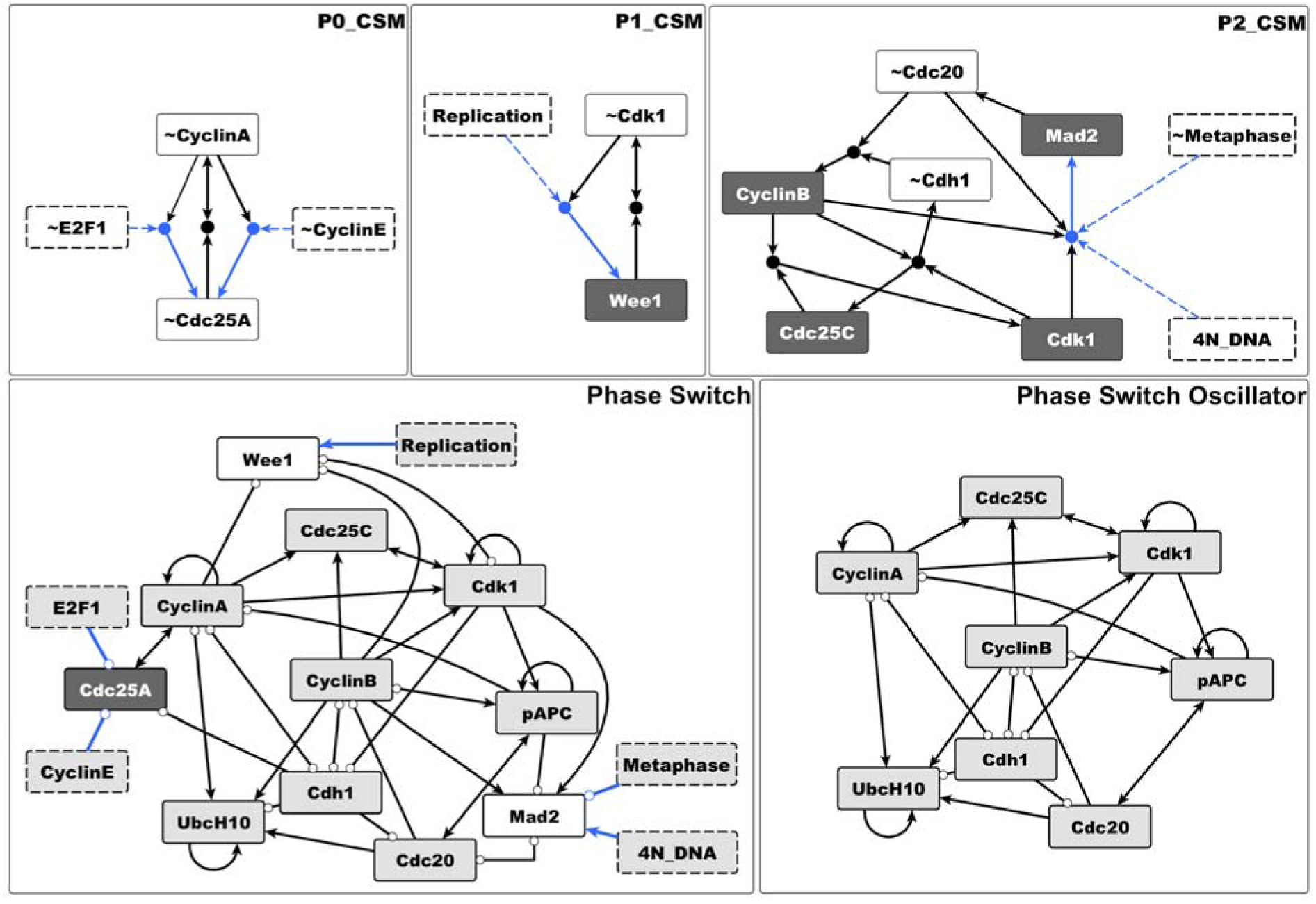
The stable motifs of the Phase Switch become conditionally stable motifs in the larger context of the cell cycle model. Due to the influences from the rest of the cell cycle network on Cdc25A, Wee1 and Mad2 (shown with blue lines), the P0 motif turns into two conditionally stable motifs which differ only in their condition (∼E2F1 or ∼CyclinE, respectively, see top left panel). The P1 motif can only stabilize if the abstract node Replication is on. P2 can only stabilize if the abstract regulator Metaphase is off and 4N DNA is on simultaneously (top right panel). The Phase Switch Oscillator (bottom right) is obtained from the Phase Switch (bottom left) by substituting the states Wee1 =Mad2=0 and Cdc25A=1. As in Figure 2, dark grey node background indicates the ON state of the node and white background refers to the OFF state.

To explore what alternative behaviors remain in the attractor repertoire of the Phase Switch, we consider an extreme scenario with respect to stabilization. Namely, we set the state of the three nodes that receive outside influences (representing the checkpoint conditions) to the state *opposite* of their states in the attractors of the Phase Switch. Namely, we substitute the values Cdc25A=1, Wee1=0 and Mad2=0 into the regulatory functions of the Phase Switch, consequently destabilizing all three of its stable motifs. The resulting system, shown in the bottom right panel of Figure 3, operates without any checkpoint control. This is akin to the network responsible for cell cycle progression in embryonic stem cells in that it doesn’t have a restriction point^47^. Contrary to the network driving embryonic stem cell division, this circuit also lacks a DNA damage and spindle assembly checkpoint.

After locking Cdc25A on, Wee1 and Mad2 off the Phase Switch module is reduced to eight nodes and 28 edges. It is still strongly connected and contains positive and negative cycles of various lengths (bottom right panel of Figure 3). Destabilizing the stable motifs of the Phase Switch might be expected to create a set of complex attractors, some of which may be dependent on the update scheme. Interestingly, we find a single limit cycle attractor of 9 states using synchronous update (Supplementary Figure S1), and a single 141-state complex attractor under asynchronous update. In recognition of this fact we refer to this modified system as the Phase Switch Oscillator (PSO). The entire state space of the PSO is one attractor basin, as all remaining states (out of the 2^8^=256 states) converge to the limit cycle / complex attractor.

### The Phase Switch Oscillator traces a robust cycle through the three attractors of the Phase Switch

To visualize the PSO’s dynamics under synchronous and asynchronous update, we sampled the most frequently visited states of the complex attractor with asynchronous update (see Methods), and overlaid the synchronous limit cycle on the resulting state transition graph (Figure 4). Consecutive states of the synchronous state transition graph differ in one to three node states (i.e. during a synchronous update one, two or three nodes change state), while the edges of the asynchronous state transition graph represent changes in a single node (indicated as edge label). Despite this difference in the definition of a state transition, the PSO’s synchronous and asynchronous attractor follow similar paths along the cell cycle. Both attractors visit the G0/G1, post-G2 and post-SAC states, and the most likely asynchronous trajectory also visits the G2 and SAC Phase Switch attractors, which are skipped in the synchronous update.

**Figure 4.**
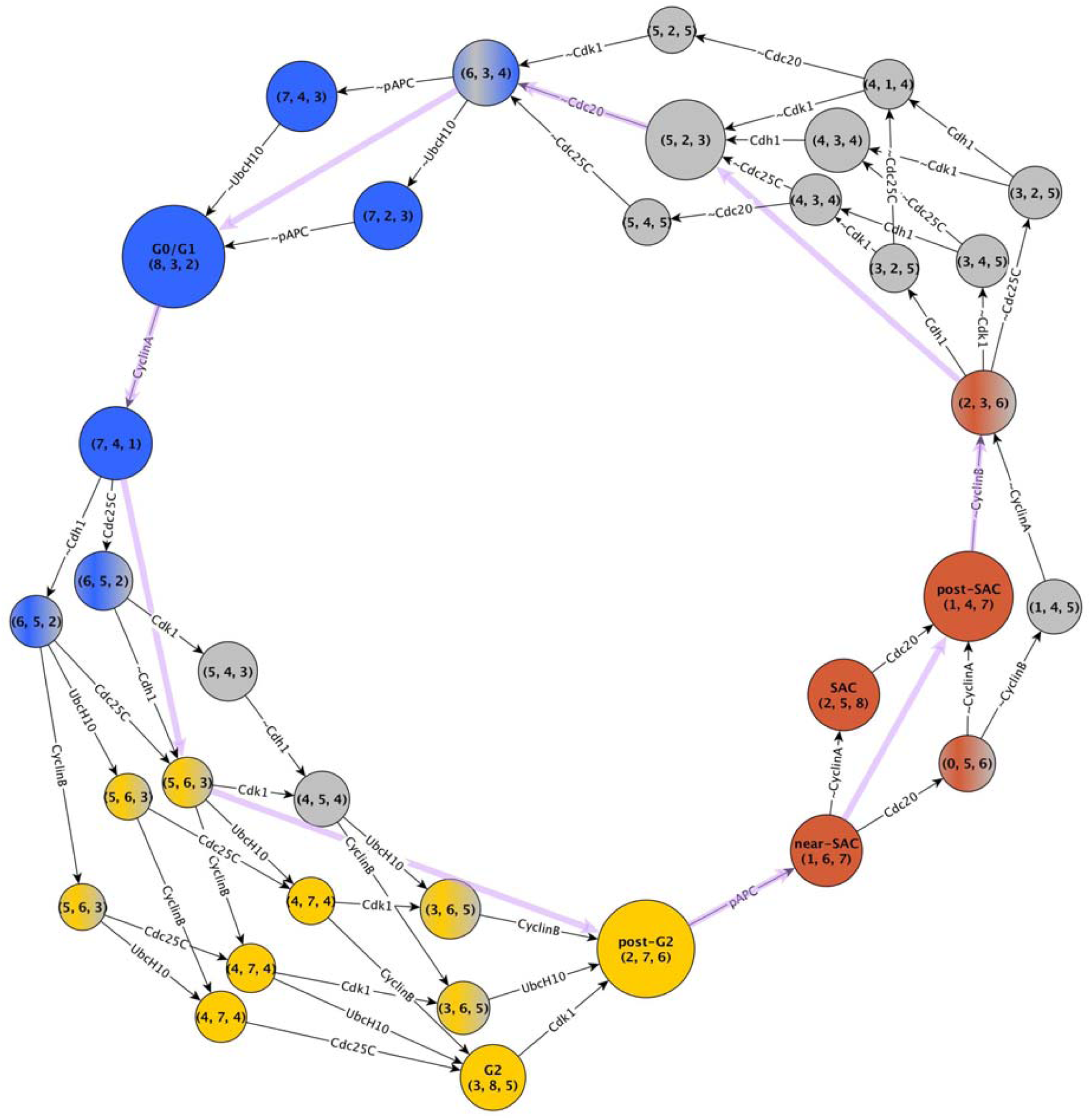
A filtered representation (see Methods) of the complex attractor of the Phase Switch Oscillator. To provide a unique identifier without indicating the state of each node, each state of the system is labeled with its overlap with the three Phase Switch attractors (in the order G0/G1, G2, SAC). If the overlap of a state with one of the tree attractors is 7 or 8 the node is colored with the color representing the relevant attractor, namely blue for G0/G1, yellow for G2 and brown for SAC. States that have an overlap of less than 6 with an attractor are shown in grey. If the overlap is 6 the state is colored with a combination of grey with the color of the respective attractor. The color combinations mirror the transitions between the phases. Some important states are marked by a label representing the closest phenotype. The node sizes represent the relative visitation probability of the corresponding states (see Methods). Each edge label shows the node that changes state in the respective transition; if the node name is preceded by ∼ the node turns off, otherwise it turns on. For simplicity we omit the self-loops that correspond to the cases where a node state is re-evaluated but does not change. The state transitions corresponding to synchronous update are shown in purple.

Supplementary Text S2 provides an edge-by-edge comparison of the two attractors, underscoring the remarkable agreement between the synchronous and asynchronous state transition graphs as both faithfully follow the cell cycle. Figure 5 compactly summarizes this agreement in a “backbone” representation of the complex attractor. The vast majority of the paths in the asynchronous attractor follow the synchronous limit cycle and robustly adhere to its temporal ordering of states (i.e., the order of phases in the cell cycle). Indeed, all the paths of the asynchronous STG are oriented in the same direction as the paths of the synchronous STG.

**Figure 5.**
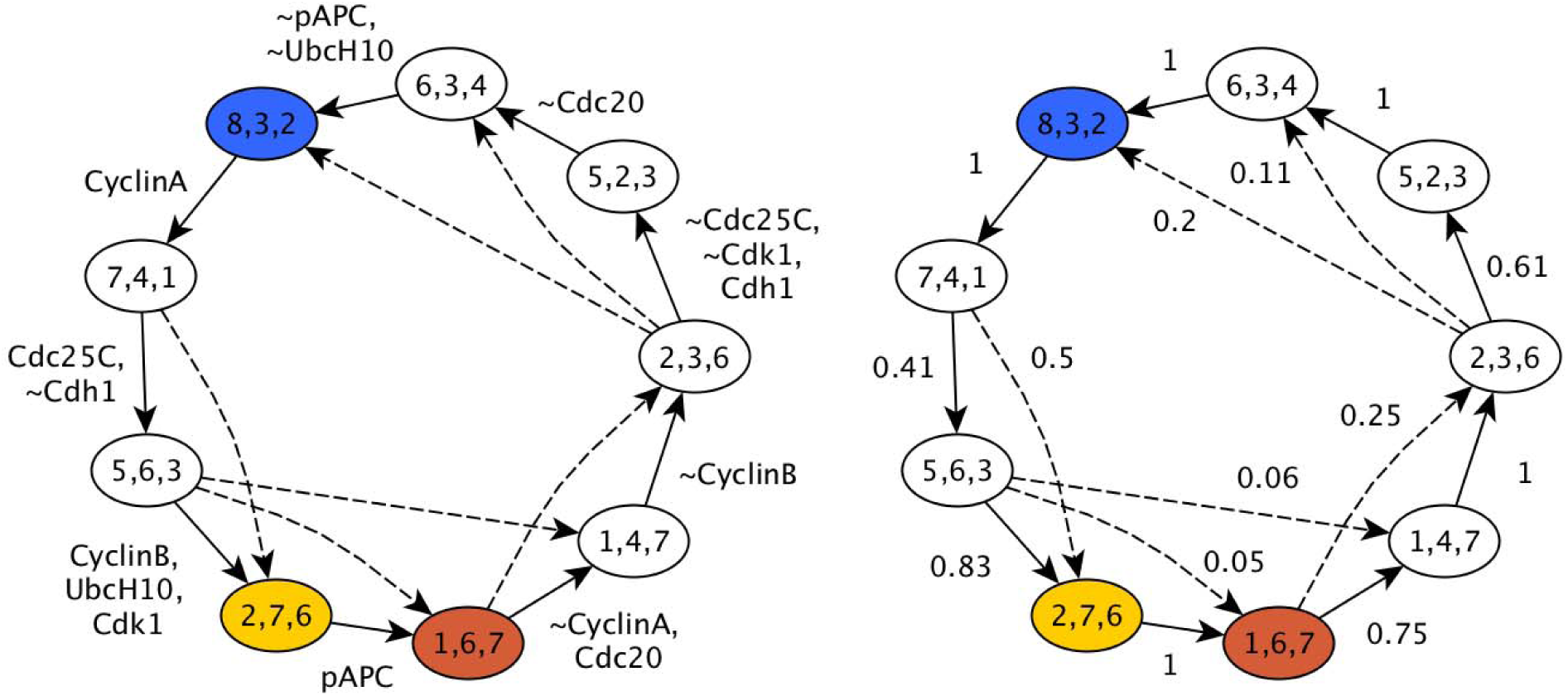
The shared backbone of the synchronous and asynchronous state transition graphs. The nodes represent the nine states visited during synchronous update; the node labels indicate the overlap of the corresponding state with the three Phase Switch attractors (in the order G0/G1, G2, SAC). The solid edges represent single state transitions obtained by synchronous update. These state transitions also appear in the asynchronous state transition graph, either as edges or as paths. The dashed edges indicate cases where paths exist in the asynchronous state transition graph that skip a state visited by the synchronous state transition graph. The states marked in blue, yellow and brown indicate the states closest to the G0/G1, G2, and SAC attractors, respectively. In the left panel the edge labels indicate the nodes that change state during the corresponding transition; nodes whose name is prefaced by ∼ turn off, the rest turn on. For each synchronous state transition the asynchronous state transition graph contains a path that corresponds to sequential state changes of the same nodes. In the right panel the labels on the state transition edges indicate the probability of the state transition when using asynchronous update. State transitions with a probability less than 0.05 are omitted from this figure. Supplementary Table S2 indicates the probability of every possible transition between these nine states.

In order to quantify how closely the asynchronous dynamics adheres to the synchronous cycle, we computed the aggregated probability of all paths of the asynchronous STG that start and end at the states of the synchronous STG without visiting other states of the synchronous STG (see Methods); these probabilities are indicated as edge labels on the right panel of Figure 5. Asynchronous update paths between limit cycle states that are not adjacent in the synchronous STG are deemed “shortcut transitions”; these transitions mix the node state changes involved in multiple steps of the synchronous update. A comprehensive list of the probability of all shortcut transitions is given in Supplementary Table S2. In spite of the random update order of the asynchronous update, only six shortcut transitions have probability higher than 0.05.

The robustness of the complex attractor is further illustrated by the distribution of the on and off durations of each node, summarized in Supplementary Figure S3. The median durations vary from 5 (for Cdc20=1, CyclinB=1 and UbcH10=0) to 11 (for CyclinB=0), but for each variable, the sum of the median on and off durations is very close to 16, i.e., the number of node state changes on the synchronous limit cycle. Similarly, the median period of oscillation for each node is very close to 16.

### The expanded network of the Phase Switch Oscillator forms an oscillating motif

To understand what causes the robustness of the oscillation and the sequential approach of the Phase Switch attractors we analyze the expanded network of the PSO, which encapsulates both the topological and logical features of the regulatory network.

The expanded network of the Phase Switch Oscillator contains 16 virtual nodes and 21 composite nodes (Supplementary Figure S4). Its 72 edges form 31 sufficient relationships between virtual nodes, each of which is either direct or mediated by a single composite node. Each of these relationships appear as a disjunctive (“or”-separated) clause in the regulatory function of the target node (Supplementary Text S1). The average in-degree of the expanded network is less than two, markedly smaller than the average in-degree of the original Phase Switch Oscillator network, which is 3.5. This illustrates that regulators need to cooperate to induce state changes in target nodes^48,49^. The whole expanded network is an oscillating motif: it is strongly connected, it is composite-closed, it contains the complementary of each virtual node, and it does not have any stable motifs. The expanded network has more than 6000 cycles (closed paths with non-repeating virtual or composite nodes), the vast majority of which are inconsistent (they contain an internal contradiction either in the virtual or composite nodes of the cycle; see Methods and Figure S4). There are 28 consistent cycles (Supplementary Figure S5) all of which have fewer than 8 nodes. All the inconsistent cycles have 8 or more nodes. This difference in cycle sizes indicates that more conditions (in terms of the state of other nodes) need to be satisfied to ensure the oscillation of a node than a sustained state of a node in the Phase Switch Oscillator. In other words, nodes must rely on each other to achieve a sustained oscillation.

To help identify nodes that play a key role in the complex attractor, we generate a weighted network of virtual nodes in which the edge weight between two virtual nodes depends on the probability that one is sufficient for the other and we determine the betweenness centrality^50^ of each node (see Methods). This analysis, summarized in Supplementary Figure S6, suggests that CyclinA, pAPC, and Cdh1 are crucial for the oscillation, as both of their virtual nodes have a near-maximal betweenness centrality. At the other extreme, the virtual nodes Cdc25C, ∼Cdc25C and ∼UbcH10 have a betweenness centrality of 0.

The expanded network embodies the logic sufficiency and necessity relationships that drive the dynamics of the system. As such, the expanded network explains the trajectories of the state transition graph (see Supplementary Text S3 for a detailed comparison of the expanded network and the complex attractor).

### The Phase Switch Oscillator contains a cycle of conditionally stable motifs that sequentially stabilize each other and cause their own destabilization

We identified all conditionally stable motifs (CSMs) in the PSO that have only a single condition (see Methods). These are depicted and labeled in Figure 6. The smallest CSM is a node that can maintain its state with the help of another node. This situation appears in the expanded network as a virtual node that is connected by a bidirectional edge to a composite node with a single additional regulator. The composite node’s other regulator serves as the condition for the CSM. There are two such nodes, pAPC and UbcH10; both virtual nodes of each form CSMs (C6, C8, C12 and C13 in Figure 6). Other elementary CSMs of the Phase Switch Oscillator contain two or three virtual nodes. Several elementary CSMs overlap in yet larger CSMs, indicating that satisfying a single condition can often stabilize relatively large subnetworks.

**Figure 6.**
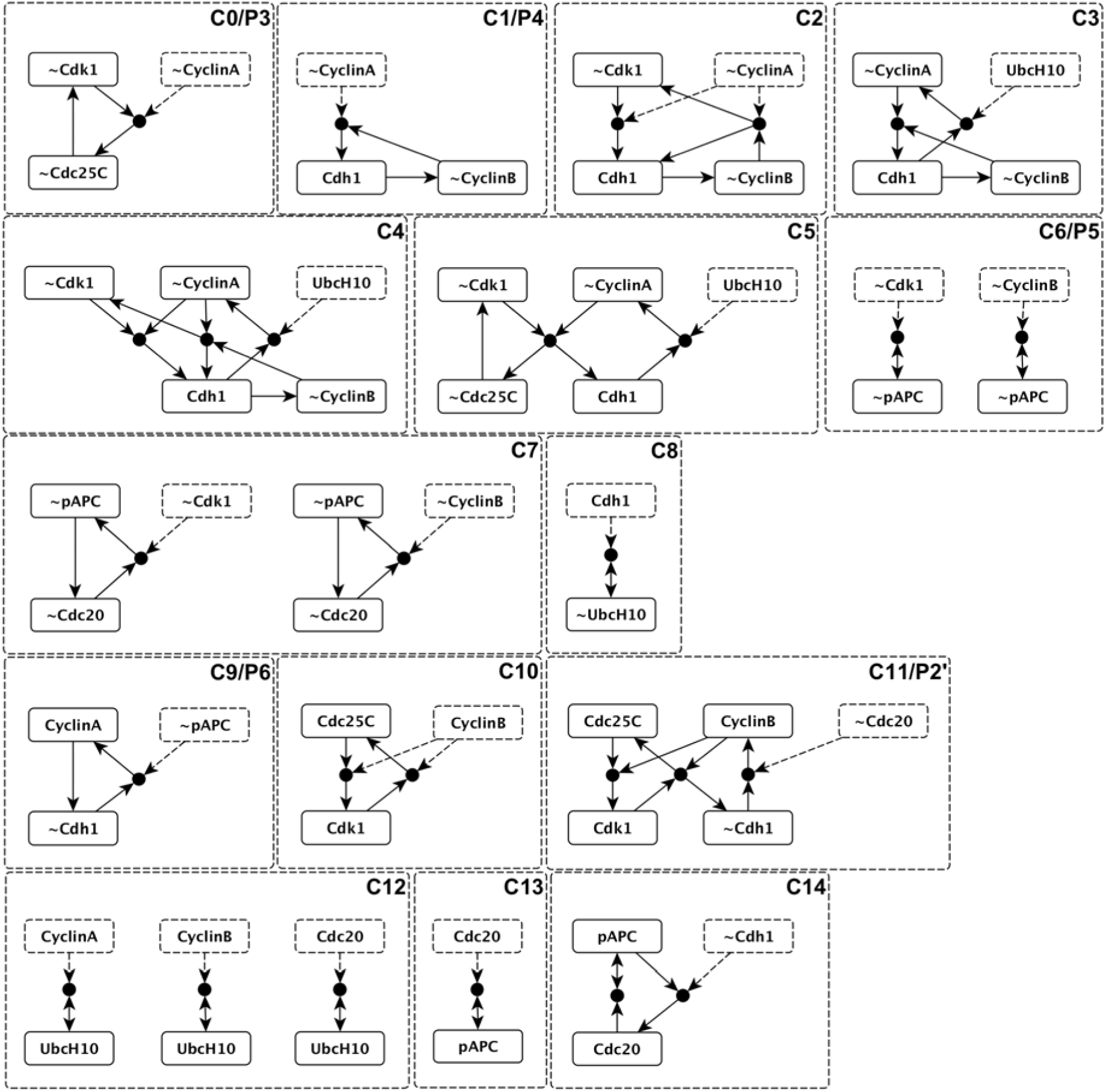
Conditionally stable motifs of the Phase Switch Oscillator (PSO). All of them are subgraphs of the expanded network of the PSO (see Supplementary Figure S4). Virtual nodes with dashed outline represent the conditions. The labels in the top right corner of the boxes indicate the name of the conditionally stable motif as well as the corresponding Phase Switch motif. Multiple motifs in the same box (e.g. C6/P5) consist of the same virtual nodes but with different conditions.

Four CSMs of the Phase Switch Oscillator coincide with the four CSMs of the Phase Switch (C0/P3, C1/P4, C6/P5, C9/P6, respectively, in Figure 6). The CSM C11 includes four virtual nodes of the P2 stable motif (thus the C11/P2’ notation). In addition, the PSO has CSMs that are not present in the Phase Switch; notably C14, which contains the Cdc20=1 state absent from all three attractors of the Phase Switch.

CSMs can be linked via their logic domains of influence. The logic domain of influence of a seed set of virtual nodes is the set of virtual nodes that are causally stabilized when the seed set is held fixed (see Methods). Using each CSM as a seed set, we identified other CSMs in its domain of influence and represented each such causal influence as a directed edge on Figure 7. In order to organize this interaction network, we delineated CSMs with boxes connected by solid arrows representing causal activation of a CSM by another. We represent the case when a virtual node of a CSM is the condition of another CSM with a dashed arrow (e.g. CyclinB, a virtual node in C11, serves as the condition of C10). In two cases a single CSM is not sufficient for the activation of the downstream one, requiring stabilization of two upstream CSMs. We represent these AND conditions with filled circles. analogous to composite nodes of an expanded network. A remarkable feature of the resulting CSM network is that the CSMs organize along a directed cycle of logic influences. As individual CSMs stabilize, they cause further stabilization of the CSMs downstream. This sequential CSM stabilization creates a robust sequence of network states, guaranteeing that the network’s dynamical trajectories are constrained regardless of stochasticity. As we will discuss later, this interaction pattern suggests a higher order logical network.

**Figure 7.**
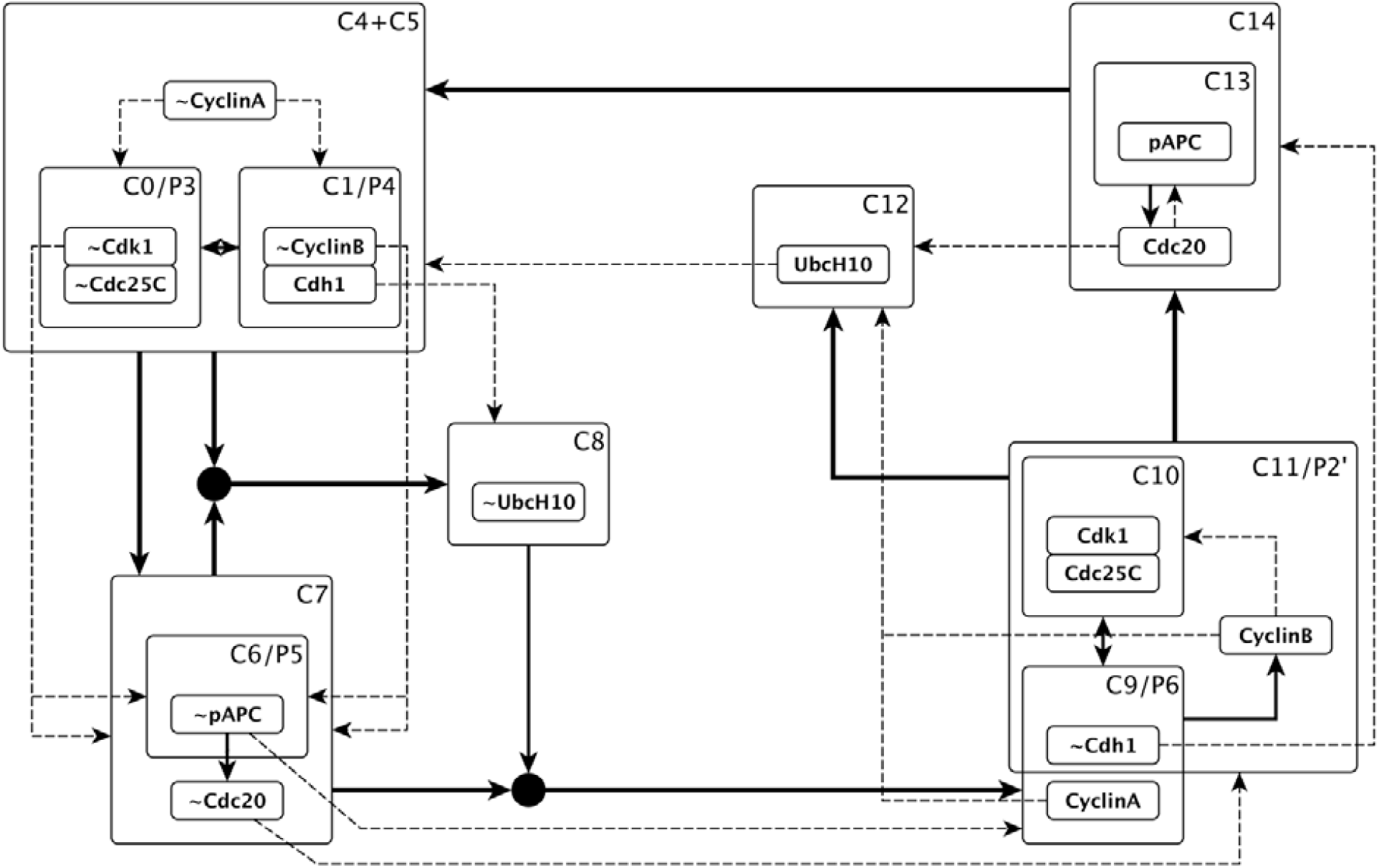
The logical relationships between conditionally stable motifs (CSMs) of the Phase Switch Oscillator. The CSMs are represented by the dashed boxes; every node state is present in at least one CSM. Dashed lines directed toward each CSM represent conditions for stability. The solid lines represent the activation of motifs, i.e. either sufficient regulators of CSMs or two regulators that together are sufficient (in which case the AND gate is represented by a black filled circle).

An additional important feature shared by many CSMs of the Phase Switch Oscillator is that they cause the deactivation of their own conditions. In other words, these CSMs have the opposite states of their own condition in their logic domain of influence. For example, the logic domain of influence of the stabilized C0 CSM, made up of the virtual nodes ∼Cdk1 and ∼Cdc25C and conditioned on ∼CyclinA, contains Cdh1, ∼CyclinB, ∼Cdc20, ∼pAPC, ∼UbcH10, and CyclinA (Supplementary Figure S7). The activation of CyclinA marks the deactivation of the condition of C0, allowing its nodes to change state. Supplementary Table S3 lists all the single-condition conditionally stable motifs that cause their own destabilization. Together, the chain of self-destabilizing CSMs on Figure 7 drive the robust oscillatory dynamics of the Phase Switch Oscillator.

### Examining the motif structure and dynamics of networks with a locked node reveals the differences between the nodes’ influence

All eight nodes of the Phase Switch Oscillator participate in the oscillation and spend a similar amount of time in their two states (see Supplementary Figure S3). All 16 virtual nodes participate in at least one CSM. We next asked whether all nodes contribute equally to the oscillation. Our betweenness centrality analysis of virtual nodes (see Supplementary Figure S6) suggests that they can be categorized in four centrality classes, the lowest being a betweenness centrality of zero. To evaluate each node state’s contribution to the complex attractor, we systematically set each node in its active or inactive state and identify the motif structure and attractor repertoire of the thus-modified dynamical system. The modified systems’ dynamic behaviors fall into three categories: 1) the PSO oscillation is preserved as the sole attractor, 2) the modified system is monostable, or 3) the modified system is multistable. The expanded networks for representatives of each of the three categories are shown in Figure 8, alongside the original system’s expanded network. A comprehensive summary of the stable and conditional motifs and attractors corresponding to each intervention is provided in Supplementary Table S4. Of the sixteen possible node-state-fixing interventions, only three, namely Cdc25c = 1, UbcH10 = 0, and Cdk1 = 1, preserve oscillation. A shared feature of these three interventions is that they are not conditions for any CSM and cannot stabilize any other node. Therefore, no stable motif is created by any of these interventions. The modified system loses the CSMs that include the complementary of the fixed state as a virtual node or as a condition (e.g. it loses motifs C4, C5 and C12 for UbcH10=0). Nevertheless, sufficiently many CSMs that induce their own destabilization remain so that the oscillation is preserved.

**Figure 8:**
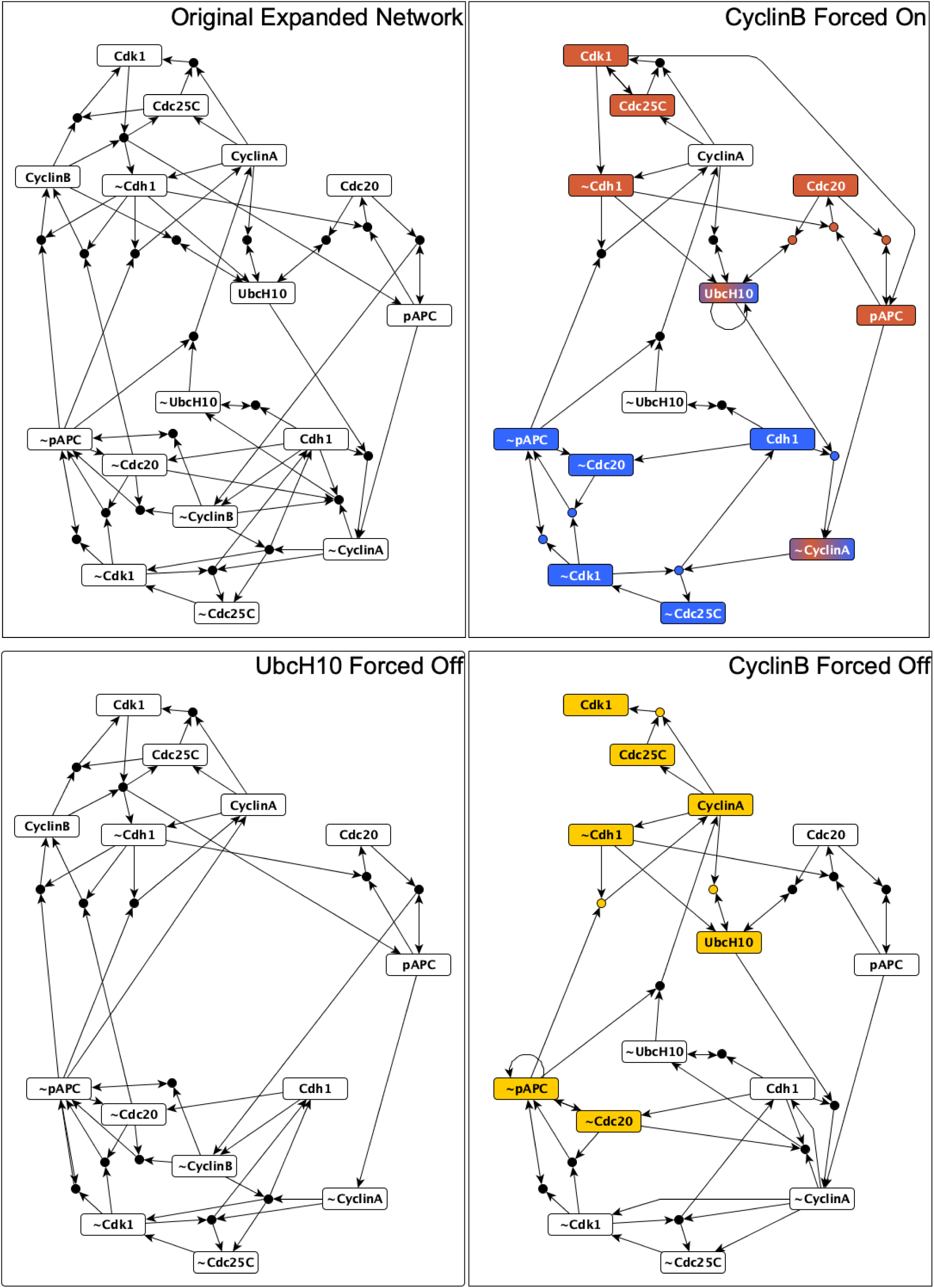
The expanded network that results from no intervention (top left panel) or three characteristic interventions (other panels as indicated by panel titles). Virtual nodes whose label is the node name preceded by ∼ indicate the off state of the relevant node. The three interventions exemplify each of the three attractor categories outlined in the main text: retained oscillation (bottom left, UbcH10 off), monostability (bottom right, CyclinB off), and multistability (top right, CyclinB on). The group of virtual nodes that make up each fixed point are highlighted in color. The blue state has the greatest overlap with the G0/G1 attractor of the Phase Switch, the yellow state most closely overlaps with the G2 attractor, and the brown state is most similar to the SAC attractor.

Eleven of the node-fixing interventions result in a reduced system that is monostable. Most newly stabilized states are highly similar to the three Phase Switch Attractors. Namely, CyclinA = 0, pAPC = 0 and UbcH10=1 each lock the PSO into a G0/G1-like state; Cdc25c = 0, Cdk1 = 0 and CyclinB = 0 each lock it into a G2-like attractor, while Cdc20 = 0 locks it into SAC. In contrast, Cdh1 = 1 stabilizes the network in a state poised to transition out of G1 into G2, CyclinA = 0 locks it poised at the boundary of G2 and mitosis (towards SAC), and Cdh1 = 0 locks it at the boundary between SAC and G0/G1. In all the cases of monostability a CSM or the union of multiple compatible CSMs becomes a stable motif, thus its influence becomes non-contradictory; and opposing CSMs disappear. In addition, two modifications result in multi-stability; Cdc20 = 1 and CyclinB = 1. Locking Cdc20 to 1 results in three qualitatively similar fixed points that only differ in the composition of the protein complex containing Cdc20 (i.e., they only differ in the values of pAPC and UbcH10); the rest of the nodes stabilize into their G0/G1 state.

In contrast, locking CyclinB on creates two highly dissimilar fixed point attractors that differ in five node states and overlap only in UbcH10 = 1 and CyclinA = 0 (see Figure 8). The bistability in the presence of forced CyclinB expression is due to the fact that sustained CyclinB is the (direct or indirect) condition for two mutually exclusive CSMs. CyclinB is the condition for the C10 CSM (Cdk1, Cdc25C), which drives an attractor resembling a cell at the point of SAC passage (brown node-states on Figure 9, top right). CyclinB is also the condition of the C12 CSM (UbcH10), which when stabilized serves as the condition for the C3 CSM (Cdh1, ∼CyclinA, ∼Cdc25C, ∼Cdk1), which when stabilized leads to an attractor resembling the G0/G1 state (blue node-states on Figure 8, top right). In contrast, when CyclinB is held inactive the C7_2 CSM (∼Cdc20, pAPC) becomes a stable motif. The activation of this motif enables the activation of the C9 CSM (CyclinA, ∼Cdh1), resulting in a single fixed point most similar to the G2 attractor. Thus, control of CyclinB can yield any of the three attractors of the Phase Switch, consistent with biological knowledge^46,51–53^.

**Figure 9.**
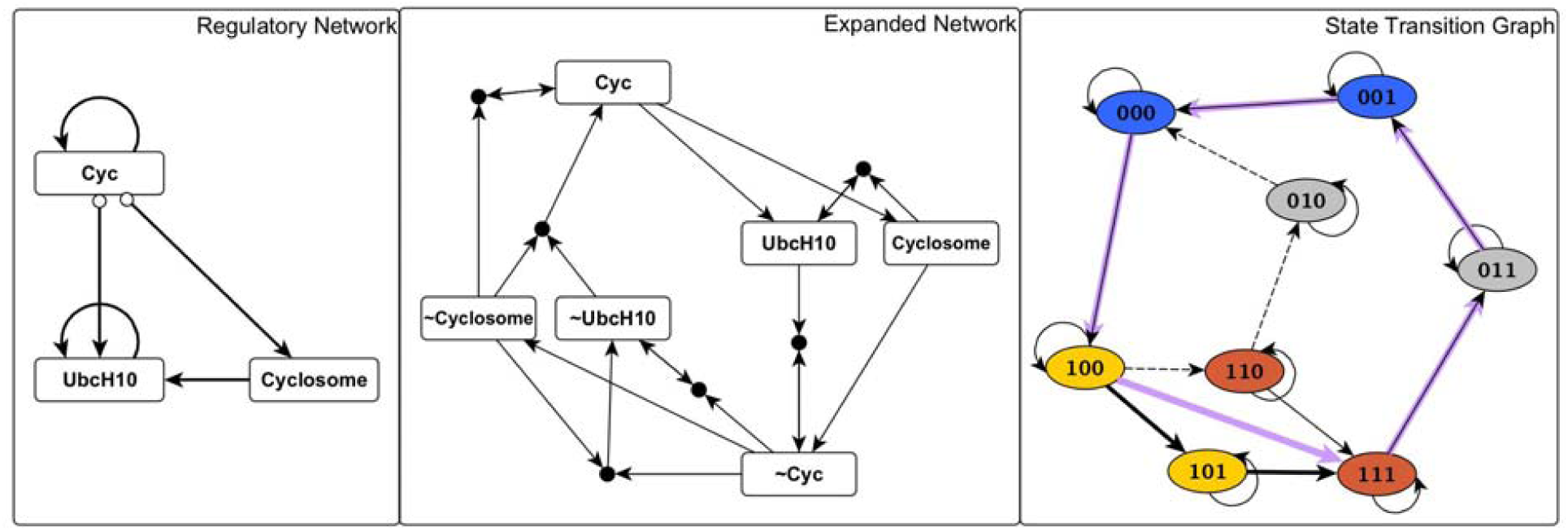
The logical relationships that determine the transitions of the PSO can be effectively illustrated by defining aggregated meta-nodes for overlapping conditionally stable motifs. The Cyc meta-node contains CyclinA, CyclinB, Cdc25C, Cdk1, and ∼Cdh1. The Cyclosome meta-node includes the virtual nodes pAPC and Cdc20. The complementary node (negation) of a meta-node includes the complementary nodes of the meta-node’s constituent virtual nodes. The first two panels indicate the regulatory and expanded network of meta-nodes. The panel on the right shows the state transition graph of the meta-node network. In this panel the labels of each state are in the order Cyc, Cyclosome, UbcH10. Node background color indicates the states closest to the attractors of the Phase Switch: blue for G0/G1, yellow for G2, brown for SAC. The synchronous cycle is shown by purple edges. The asynchronous complex attractor is made up by two cycles of unequal size, but each of which approaches the three attractors in the same order. The difference between the two cycles is whether UbcH10 activates and then deactivates (longer cycle) or stays inactive (short cycle). This latter process has a very low probability.

In addition to stabilizing any of the three Phase Switch attractors when locked on or off, step-wise changes between fixed states of CyclinB can induce attractor transitions that mimic cell cycle progression (Supplementary Figure S8). Indeed, the attractor of the CyclinB = 0 constrained Phase Switch Oscillator is in the basin of attraction of the SAC-like attractor of the CyclinB = 1 constrained PSO (in particular, the C10 conditionally stable motif is active). Thus, if CyclinB is absent for a long duration, and is then reintroduced, the model system undergoes a transition from a G2-like state to a SAC-like state; a model behavior that matches experimental observations^51–53^. Furthermore, if we artificially remove CyclinB *after* the SAC-like state is reached in the CyclinB = 1 constrained system, then with high probability, the system passes through the neighborhood of the G0/G1 state, which was not visited by the trajectory from the G2-like state to the SAC-like state (Supplementary Figure S8). Reintroducing CyclinB when the system is in this region of state space drives it into the G0/G1-like attractor^54–56^. A similar hysteresis in response to the increase vs. decrease of CyclinB was experimentally observed in Xenopus embryonic cells^52^. In conclusion, we predict that there exists a sequence of repeated changes in CyclinB that can drive the system to visit the attractors of the Phase Switch in the same order as the cell cycle.

### A higher-level network of three nodes qualitatively replicates the oscillation

The interplay of the conditionally stable motifs allows for a concise, coarse-grained description of the complex attractor. Inspection of Figure 7 suggests a grouping of multiple overlapping CSMs into larger CSMs. One large group unites conditionally stable motifs C0-C5 and is composed of the virtual nodes ∼Cdk1, ∼Cdc25C, ∼CyclinB, Cdh1, and ∼CyclinA. Strikingly, these virtual nodes’ complementary nodes also form a overlapping group of conditionally stable motifs, namely C9-C11. These two groups can thus be represented by complementary meta-nodes. We define the meta-node as being formed by CyclinA, CyclinB, Cdc25C, Cdk1, ∼Cdh1, and name it “Cyc”. Similarly, the virtual nodes pAPC and Cdc20 (as part of CSMs C13, C14) form a group that can be represented by a meta-node (which we denote “Cyclosome”), and ∼pAPC and ∼Cdc20 (as part of CSM C7) form the complementary meta-node. Together with the two states of UbcH10, these four meta-nodes correspond to the largest CSMs of the expanded network (see Methods). With these notations, the higher order logic of the CSM meta-nodes can be distilled into the regulatory functions

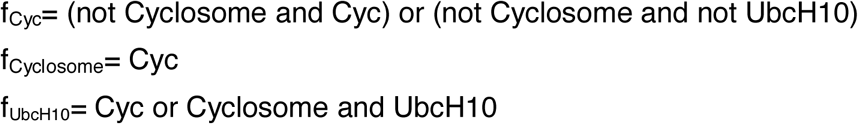

These regulatory functions express the following logical relationships: Cyclosome and UbcH10 inactivate Cyc; the only possibility for Cyc activation is the simultaneous inactivity of both Cyclosome and UbcH10. Existing Cyc activity can be maintained if Cyclosome is inactive. Cyc is sufficient for the activation of Cyclosome and UbcH10. UbcH10 activity can also be sustained if Cyclosome is active. Figure 9 illustrates the regulatory and expanded networks of meta-nodes as well as the corresponding STG.

The higher-order regulatory logic of the CSM meta-nodes explains the PSO’s cycling between states close to the G0/G1, G2, SAC attractors. The G0/G1 state corresponds to the inactivity of all three meta-nodes. This is not an attractor; inactivity of pAPC and Cdc20 implies that Cyc will activate. When this is achieved, the system is closest to the G2 attractor of the original Phase Switch. Next, the activity of Cyc drives the activation of UbcH10 and Cyclosome in an order determined by the relative speed of establishing conditionally stable motif C12 versus C14. The probability of UbcH10 turning on before Cdc20 and pAPC is at least 0.986 (see Supplementary Table S2). The rare case in which Cyclosome activates before UbcH10 leads to a path of state transitions from the G2 state to the G0/G1 state, through a SAC-like state that has UbcH10=0, consistent with the observation that fixing UbcH10 off preserves the complex attractor. In the more likely scenario, UbcH10 activates first, and then the activation of the Cyclosome node (pAPC and Cdc20) marks the Spindle Assembly Checkpoint. This state is also short-lived, as Cyc is deactivated by Cyclosome, which in turn causes the inactivation of both Cyclosome and UbcH10. Thus, the system returns to the G0/G1 state.

This reduced model supports the idea that the interplay of the CSMs is responsible for the robust cyclic behavior of the PSO and its approach to the fixed-point attractors of the Phase Switch model (see Supplementary Table S5). We argue that this is possible because all CSMs of the Phase Switch remain intact in the PSO, including the remainder of the P2 stable motif after its Mad2 node is locked off, which turns into the conditionally stable motif P2’. Their activation inevitably drives the PSO towards the attractors of the Phase Switch, thus preserving the functionality of the original module.

## Discussion

The activation of stable motifs in multi-stable Boolean networks mark points of no-return in the trajectory towards a fixed point. Here we introduced the related concept of conditionally stable motifs, which can maintain a fixed state of their constituent nodes as long as the state of one or more nodes is maintained. Through our analysis of the Phase Switch and the Phase Switch Oscillator we have uncovered three key features of CSMs. First, they can play an important role in the decision-making of multi-stable systems. For example, in the Phase Switch model the primary stable motif P1 is compatible with both G0/G1 and G2 attractors. The subsequent stabilization of the conditionally stable motif P5 and P0 or, alternatively, P5 and P6, steers the system into one or the other attractor respectively (see Figure 2). Second, in systems with complex attractors, CSMs reduce noise introduced by stochastic update order by temporarily eliminating degrees of freedom. Indeed, CSMs with the fewest conditions capture the temporary stability of short-range positive feedback loops, which temporarily fix the states of the nodes in the feedback loop. Third, oscillation requires that no CSM has stabilized conditions, and therefore the pattern of CSM condition activation and deactivation can illuminate the nature of the oscillation. In particular, when CSMs are overlapping and nested, as in the Phase Switch Oscillator, the patterns of conditional activation and deactivation for the largest CSMs yields an insightful coarse-grained description. The six states of the UbcH10, Cyc, and Cyclosome meta-nodes in states of the coarse-grained PSO correspond to the largest CSMs of the PSO. As indicated in Figure 9, the coarse-grained network reveals the long-range negative feedback loops of the Phase Switch Oscillator.

Stable motifs are well defined within the context of a model. Yet, as all models are ultimately incomplete, it is always possible that a more complete model would have additional regulators that transform the stable motif into a conditionally stable motif. In the larger context of the cell cycle model the nodes E2F1 and CyclinE of the Restriction Switch and the abstract nodes Replication, Metaphase and 4N DNA regulate nodes of the Phase Switch. Because of this external regulation the motifs P0-P2, which are stable motifs in the isolated Phase Switch, become conditionally stable motifs. We focused our analysis on an extreme scenario with respect to stabilization of the attractors of the Phase Switch, i.e. the fixed states Mad2 = Wee1 = 0, Cdc25A = 1, which represent a cell that is lacking the restriction, DNA damage and spindle assembly checkpoints. Each of these states contradicts one of the three stable motifs of the Phase Switch. The virtual nodes corresponding to these fixed node states have out-edges only to composite nodes, thus their logic domain of influence is empty, meaning that no further node states stabilize as a direct consequence of their stabilization. Moreover, these fixed node states do not create new stable motifs. The absence of stable motifs implies that the Phase Switch Oscillator does not have fixed point attractors. We hypothesize that the general condition that the fixed state of a node does not create new stable motifs is that there are no conditionally stable motifs conditioned on this fixed state. In the Phase Switch there is only one alternative combination of fixed node states that eliminates the original stable motifs and does not create new ones: Cdc25A=1, Mad2=0, Cdk1=1. This alternative combination has the same biological meaning as the one we considered.

Our analysis yields novel biological insights and predictions. For example, in contrast to the Phase Switch where Cdc20 is off in all three attractors, the emergent cycle of the PSO highlights the role of the active Cdc20 in closing the cycle by ending mitosis and driving the cell back into the G0/G1 state. Our analysis of the Phase Switch Oscillator with a locked-in state of CyclinB confirms the important role of CyclinB in driving the cell cycle of embryonic cells, and mitosis in somatic cells. We predict that there exists a sequence of repeated changes in CyclinB that can drive the system to visit the attractors of the Phase Switch in the same order as the cell cycle (see Supplementary Figure S8). More broadly, our findings support the conclusion that a combination of bistability and negative feedback underlies many biochemical oscillators^27,52^. Our newly introduced concept of conditionally stable motif may also be helpful in addressing biological learning and adaptation in a network framework^30^.

Construction of expanded networks by integrating the regulatory logic into each network was a necessary step toward identifying the possible dynamic outcomes of the interplay between positive and negative feedback loops. Both the Phase Switch and Phase Switch Oscillator networks are strongly connected and they both contain multiple negative feedback loops (Figure 3 bottom panel). Yet, neither system’s expanded network (Supplementary Figures S2 and S4) has any paths through which a single virtual node can induce its own negation. At least two virtual nodes together are required to induce the negation of one of them. For example, CyclinA and CyclinB together can stabilize the states that make up the Cyc meta-node, which leads to the activation of the Cyclosome meta-node, which yields the deactivation of CyclinA and CyclinB (accompanied by the reversal of all states in the Cyc meta-node, see Figure 9). In other words, the negative feedback loops are not functional in isolation. Both the Phase Switch and Phase Switch Oscillator contain strongly connected subgraphs that lack negative feedback loops (in other words, they are sign-consistent). The largest such subgraph of the Phase Switch contains eight nodes (all but pAPC, Cdc20, UbcH10). The largest such subgraph of the Phase Switch Oscillator coincides with the Cyc meta-node, and the second largest is the Cyclosome meta-node. While in the case of the Phase Switch Oscillator it would have been possible to define a coarse-graining based on the structure alone (especially with the benefit of hindsight), no structure-based coarse-graining can explain the attractors of the Phase Switch because they involve different subsets of the eight-node subgraph.

The expanded network framework is part of a broader effort to characterize and represent as a network the causal relationships between variables of a dynamical system. Related concepts include signed interaction hypergraphs^21^ and dynamics canalization maps^48^. The logic domain of influence of a node state is conceptually similar to the three-valued (0, 1, unknown) logical steady state that results from fixing a node state^21^ and to the dynamical modules of dynamics canalization maps, which represent the states inexorably stabilized by an input configuration^48^. The concepts of expanded network and stable motif have been generalized and implemented in multi-level discrete systems and continuous-variable systems described by ordinary differential equations^32,44^. When considered generally, the expanded network encodes causal links between regions of state-space. Each of its virtual nodes represents a region of state-space (e.g., the region in which a particular variable takes a specified value or range of values) and the composite nodes represent the intersection of the virtual node regions. Once an expanded network is constructed for a given dynamical system, be it discrete or continuous, the concept of a conditionally stable motif is immediately applicable. Thus, it is possible that by using the methods of Rozum & Albert 2018^44^, many of the concepts we have introduced here can be generalized to multi-level discrete dynamical systems and ODE models.

Our analysis indicates that the influences on the Phase Switch originating from the other modules become functional in a manner that allows the Phase Switch to approach one of its attractors (as the cell reaches the relevant checkpoint), but then they destabilize this attractor as the checkpoint is cleared. In other words, the network around the Phase Switch helps provide the *conditions* that govern its stable motifs. Nevertheless, we found that the robust channeling of its dynamics along a limit cycle is intrinsic to this network. Thus, the Phase Switch balances the need to stably maintain the cell at each checkpoint with the need for a robust limit cycle when checkpoints are cleared without issue. The methodology described in this paper can be used to understand the complex oscillation that emerges from the coupling of the Phase Switch and the Restriction Switch in the presence of growth factors. As explained in Deritei et al. 2016^17^, this attractor faithfully reproduces the cell cycle in the presence of the checkpoints we removed in this study, while toggling the combinations of the module attractors. Analyzing the conditionally stable motifs of the coupled model could shed light on an even more comprehensive coarse-grained logical network that drives the cell cycle, and offer further insight on dynamical modularity.

## Methods

### Mapping the complex attractor

The trajectories generated by general asynchronous update can be interpreted as a random walk on the state transition graph. During a random walk on a directed network a walker on node i at time step t randomly chooses one of the edges going out of i and traverses that edge in one time step. Using general asynchronous update every node has the same probability of being updated, thus every outgoing edge (i.e. every state transition that yields a different state than the starting state) has the same probability. An extended simulation of the model trajectory starting from any node in the basin of attraction of the complex attractor (i.e. the states that are starting points of trajectories that reach the complex attractor, which in the case of the PSO is the entire state space) gives us a reliable sample of the most probable states and transitions. We performed a random walk of 100,000 steps and determined the visitation counts of states and transitions. We then filtered the state transition graph, leaving only nodes that were visited at least 500 times. We validated the visitation probabilities emerging from our sampling process by using the PageRank algorithm^57^ on the full complex attractor. Mapping of the full state transition graph of the eight-node PSO is tractable; this would not be the case for larger networks.

### Coarse-graining the complex attractor based on established proxy states

On the state transition graph corresponding to general asynchronous update each outgoing edge has the same probability (as all node updates are equiprobable). We define the edge probability as the inverse of the out-degree of the edge’s source node. Using the complete general asynchronous state transition graph, we determine the transition probability between the nine states of the synchronous attractor, which we termed proxy states. For each pair of proxy states, we find all simple paths between the two states that do not involve any other proxy state and calculate the probability of each path as the product of edge probabilities over the path. The transition probability between the two proxy states is the sum of the path probabilities. We use these probabilities to construct the backbone of the complex attractor: starting with a disconnected graph of proxy states, we consider each pair of proxy states, determine the transition probability between them, and add an edge between them if the transition probability is non-zero.

### Identification of Conditionally Stable Motifs

Every CSM is a strongly connected component of the expanded network, and therefore it is either a cycle or can be viewed as a union of directed cycles. The latter follows from the fact that every node in a strongly connected component must have a path that leads to itself, and for every edge (i,j) in the component, there is a path from node j to node i. To satisfy the consistency criterion of the CSM definition, the corresponding cycle-set must be consistent in the sense that no virtual node or composite node in any of the cycles may be logically incompatible with any other node of any cycle in the set. Furthermore, the cycles must collectively form a strongly connected component, which precludes a set of disjoint cycles. The cycle sets subject to these two criteria (consistency and lack of a disjoint partition) are in one-to-one correspondence with the CSMs in the expanded network.

We leverage this fact to enumerate all CSMs. We begin by considering every self-consistent cycle in the expanded network. This is equivalent to considering every positive feedback loop in the regulatory network^31^. We treat these as nodes in a cycle graph. Undirected edges are drawn between a pair of nodes in the cycle graph if the corresponding cycles share a node in the expanded network and are mutually consistent (see Supplementary Figure S8). If two cycles are connected, then there is a path from any node in one cycle to any node of the other, and so the union of the corresponding cycles is strongly connected. In the same vein, any connected subgraph of the cycle graph corresponds to a strongly connected component of the expanded network. If a subgraph in the cycle graph is not connected, then its components constitute a disjoint partition of the union of cycles. Therefore, every CSM node set corresponds to either a single cycle or a connected subgraph of the cycle graph. The reverse is not necessarily true: connected subgraphs of the cycle graph might not be composed of consistent cycles. Consistent connected subgraphs of the cycle graph, however, are in one-to-one correspondence with the CSMs of the expanded network.

In the PSO, the number of positive feedback loops is small enough that the connected subgraphs of the cycle graph can be exhaustively constructed (Supplementary Figure S9). The computational complexity of our current implementation scales combinatorially in the number of positive feedback loops, which itself can increase dramatically with the number of nodes in the network. Development of more efficient or heuristic methods will be the focus of future work.

### Characterizing node contribution to the complex attractor

We consider a network composed of virtual nodes (one virtual node for each state of the network nodes). We place a weighted edge from each virtual node *u* (e.g., Cdk1=1) to each other virtual node *v* as follows. First, if the variable corresponding to *u* does not regulate the variable corresponding to *v*, no edge is placed. Otherwise, we calculate the sufficiency probability *s(u, v)* that u is sufficient for v (e.g. Cdk1=1 is sufficient for Cdh1=0), assuming that all other nodes take each of their two values with 50% probability. We assign an edge weight of 1/*s(u, v)* for each ordered pair (*u,v).* Intuitively, the higher the sufficiency of an edge, the closer the parent node is to the child node, which is represented as a lower edge weight. As a concrete example, let *u* be Cdk1=1 and *v* be pAPC=1. The regulatory function for pAPC is f_pAPC_= CyclinB and Cdk1 or pAPC and Cdc20.

Cdk1=1 is sufficient for pAPC=1 if and only if CyclinB is active, which we assume occurs with 50% probability. Therefore, *s(u, v)* is 0.5, and so we assign a weight of 2 to the edge from Cdk1=1 to pAPC=1.

After this network is constructed, we compute the betweenness centrality score for each virtual node. The betweenness centrality of a node, *i*, is the sum over all node pairs (*j,k*) of the fraction of shortest paths from *j* to *k* that pass through *i*^*50*^. As frequently done, we divide this score by the highest observed score, such that the values on Supplementary Figure S6 are between 0 and 1. The relative scores estimate the likelihood that the associated node state will be achieved as information spreads through the network from one randomly selected node to another. We view this as a proxy for the importance of a node in sustaining the observed oscillation.

### Logic domain of influence

For a given “seed” set of virtual nodes, one can define a logic domain of influence (LDOI) subgraph on the expanded network^58^. Informally, the LDOI of a seed set of virtual nodes is the set of node states that are causally stabilized by the seed set when it is held fixed. We used the LDOI identification algorithm developed in Yang et al. 2018, available at https://github.com/yanggangthu/BooleanDOI. In this algorithm^58^ the LDOI is defined and built via an iterative process, beginning with the empty set. On each iteration, every child node of every node in the seed set is considered (breadth first) and added to the LDOI set if the child node does not contradict any nodes in the seed set and either 1) the child node is a virtual node or 2) the child node is a composite node and all its parent nodes are already in the set. This process is continued, considering child nodes of the nodes already included in the LDOI, until no new nodes can be added. The LDOI of a stable motif includes the motif itself and no contradictions^58^. Here, we modify this definition slightly in order to better describe the PSO. When a child virtual node contradicts nodes in the seed set, we include the child node in the LDOI, but do not include any of its children. In Yang et al. 2018^58^ it is proven that all contradictions arising during the iterative process involve a node in the seed set. If the LDOI of a set contains a node that contradicts the seed set, then there is no attractor in which all node states represented in the seed set are stabilized. A corollary of this is that if the LDOI of a conditionally stable motif contains a virtual node that contradicts the condition of the CSM, the CSM can never stabilize.

### Implementations

For synchronous and asynchronous simulation of the dynamic models of the Phase Switch and Phase Switch Oscillator we used the BooleanNet python library available at https://github.com/ialbert/booleannet

The identification of the stable motifs was done using the Java library available at https://github.com/jgtz/StableMotifs

The building of the expanded network based on the regulatory functions was done using the BooleanDOI python library available at https://github.com/yanggangthu/BooleanDOI

A descriptive supplementary Jupyter Notebook that reproduces our main computational results is available at: https://github.com/deriteidavid

The notebook includes the following implementations (see Methods for details):

1. creating a BooleanNet instance of the PSO model
2. identifying the synchronous attractor by simulation
3. sampling the complex attractor with general asynchronous update scheme, where the resulting STG can be arbitrarily filtered and exported into a graphml object. This includes the comparison of the states with the Phase Switch attractors.
4. determining the full STG of the PSO and using the PageRank algorithm to validate the filtered sample
5. determining the coarse grained ‘backbone’ of the complex attractor
6. generating the expanded network of the PSO
7. finding the conditionally stable motifs of the PSO

## Acknowledgements

This work was supported by NSF grants PHY 1545832, MCB 1715826, IIS-1814405 to R.A. E.R.R was partially supported by NIH/NHLBI grant HL119322. The authors thank Jorge G. T. Zanudo, Xiao Gan, Parul Maheshwari, and David Wooten for insightful discussions.

## Author contributions statement

D.D. and R.A. conceived the study. D.D., J.R and R.A. performed the analysis. D.D. and J.R. developed and implemented the algorithms. D.D., J.R. and R.A. created the visualizations. E.R.R. validated the biological implications of the analysis. D.D., R.A., J.R. and E.R.R. wrote the paper.

## Supplementary Information

Supplementary Text S1, Regulatory functions of the Phase Switch and Phase Switch Oscillator.

Supplementary Text S2, Detailed description of the complex attractor of the Phase Switch Oscillator

Supplementary Text S3, Detailed description of the agreement of the expanded network and complex attractor of the Phase Switch Oscillator

Supplementary Figure S1, The synchronous attractor of the Phase Switch Oscillator.

Supplementary Figure S2, The expanded network of the Phase Switch.

Supplementary Figure S3, The distribution of the duration of the sustained on or off state Supplementary of each node on the complex attractor of the Phase Switch Oscillator

Supplementary Figure S4, The expanded network of the Phase Switch Oscillator embodies the logic relationships that drive its oscillating behavior

Supplementary Figure S5, The length distribution of consistent and inconsistent cycles on the expanded network of the Phase Switch.

Supplementary Figure S6, Betweenness centrality analysis of the expanded network of the Phase Switch Oscillator.

Supplementary Figure S7, Illustration of a conditionally stable motif causing its own destabilization.

Supplementary Figure S8, A sequence of sustained states of CyclinB can induce a sequence of transitions between attractors that mimics the cell cycle progression.

Supplementary Figure S9, The cycle graph constructed for the Phase Switch Oscillator.

Supplementary Table S1, The attractors of the Phase Switch and their biological explanation

Supplementary Table S2, Probability of state transitions among pairs of states of the synchronous limit cycle in the complex attractor found by general asynchronous update.

Supplementary Table S3, Self-destabilizing conditionally stable motifs

Supplementary Table S4, Node state intervention table

Supplementary Table S5, Phase Switch Oscillator states closest to Phase Switch attractors.

## References

1. Grieco, L. et al. Integrative Modelling of the Influence of MAPK Network on Cancer Cell Fate Decision. PLoS Computational Biology 9, e1003286 (2013).

2. Gong, H. Analysis of intercellular signal transduction in the tumor microenvironment. BMC Syst. Biol. 7 Suppl 3, S5 (2013).

3. Helikar, T., Konvalina, J., Heidel, J. & Rogers, J. A. Emergent decision-making in biological signal transduction networks. Proceedings of the National Academy of Sciences 105, 1913–1918 (2008).

4. Verlingue, L. et al. A comprehensive approach to the molecular determinants of lifespan using a Boolean model of geroconversion. Aging Cell 15, 1018–1026 (2016).

5. Zhang, R. et al. Network model of survival signaling in large granular lymphocyte leukemia. Proc. Natl. Acad. Sci. U. S. A. 105, 16308–16313 (2008).

6. Novák, B. & Tyson, J. J. A model for restriction point control of the mammalian cell cycle. Journal of Theoretical Biology 230, 563–579 (2004).

7. Toettcher, J. E. et al. Distinct mechanisms act in concert to mediate cell cycle arrest. Proceedings of the National Academy of Sciences 106, 785–790 (2009).

8. Gerard, C. & Goldbeter, A. Temporal self-organization of the cyclin/Cdk network driving the mammalian cell cycle. Proceedings of the National Academy of Sciences 106, 21643–21648 (2009).

9. Albert, R. & Othmer, H. G. The topology of the regulatory interactions predicts the expression pattern of the segment polarity genes in Drosophila melanogaster. Journal of Theoretical Biology 223, 1–18 (2003).

10. Li, F., Long, T., Lu, Y., Ouyang, Q. & Tang, C. The yeast cell-cycle network is robustly designed. Proceedings of the National Academy of Sciences 101, 4781–4786 (2004).

11. Espinosa-Soto, C. A Gene Regulatory Network Model for Cell-Fate Determination during Arabidopsis thaliana Flower Development That Is Robust and Recovers Experimental Gene Expression Profiles. THE PLANT CELL ONLINE 16, 2923–2939 (2004).

12. Faure, A., Naldi, A., Chaouiya, C. & Thieffry, D. Dynamical analysis of a generic Boolean model for the control of the mammalian cell cycle. Bioinformatics 22, e124–e131 (2006).

13. Davidich, M. I. & Bornholdt, S. Boolean Network Model Predicts Cell Cycle Sequence of Fission Yeast. PLoS ONE 3, e1672 (2008).

14. Schlatter, R. et al. ON/OFF and Beyond - A Boolean Model of Apoptosis. PLoS Computational Biology 5, e1000595 (2009).

15. Calzone, L. et al. Mathematical Modelling of Cell-Fate Decision in Response to Death Receptor Engagement. PLoS Computational Biology 6, e1000702 (2010).

16. Steinway, S. N. et al. Network Modeling of TGF Signaling in Hepatocellular Carcinoma Epithelial-to-Mesenchymal Transition Reveals Joint Sonic Hedgehog and Wnt Pathway Activation. Cancer Research 74, 5963–5977 (2014).

17. Deritei, D., Aird, W. C., Ercsey-Ravasz, M. & Regan, E. R. Principles of dynamical modularity in biological regulatory networks. Scientific Reports 6, (2016).

18. Albert, R. et al. A new discrete dynamic model of ABA-induced stomatal closure predicts key feedback loops. PLOS Biology 15, e2003451 (2017).

19. Sizek, H., Hamel, A., Deritei, D., Campbell, S. & Regan, E. R. Boolean model of growth signaling, cell cycle and apoptosis predicts the molecular mechanism of aberrant cell cycle progression driven by hyperactive PI3K. PLOS Computational Biology 15, e1006402 (2019).

20. Thomas, R. & D’Ari, R. Biological Feedback. (CRC Press, 1990).

21. Klamt, S., Saez-Rodriguez, J., Lindquist, J. A., Simeoni, L. & Gilles, E. D. A methodology for the structural and functional analysis of signaling and regulatory networks. BMC Bioinformatics 7, 56 (2006).

22. Paulevé, L., Magnin, M. & Roux, O. Static analysis of Biological Regulatory Networks dynamics using abstract interpretation. Mathematical Structures in Computer Science 22, 651–685 (2012).

23. Mochizuki, A., Fiedler, B., Kurosawa, G. & Saito, D. Dynamics and control at feedback vertex sets. II: A faithful monitor to determine the diversity of molecular activities in regulatory networks. Journal of Theoretical Biology 335, 130–146 (2013).

24. Angeli, D., Ferrell, J. E. & Sontag, E. D. Detection of multistability, bifurcations, and hysteresis in a large class of biological positive-feedback systems. Proceedings of the National Academy of Sciences 101, 1822–1827 (2004).

25. Craciun, G. & Feinberg, M. Multiple Equilibria in Complex Chemical Reaction Networks: II. The Species-Reaction Graph. SIAM Journal on Applied Mathematics 66, 1321–1338 (2006).

26. Remy, É., Ruet, P. & Thieffry, D. Graphic requirements for multistability and attractive cycles in a Boolean dynamical framework. Advances in Applied Mathematics 41, 335–350 (2008).

27. Novák, B. & Tyson, J. J. Design principles of biochemical oscillators. Nature Reviews Molecular Cell Biology 9, 981–991 (2008).

28. Ingolia, N. T. Topology and robustness in the Drosophila segment polarity network. PLoS Biol. 2, e123 (2004).

29. Maheshwari, P. & Albert, R. A framework to find the logic backbone of a biological network. BMC Systems Biology 11, (2017).

30. Csermely, P. The Wisdom of Networks: A General Adaptation and Learning Mechanism of Complex Systems. BioEssays 40, 1700150 (2018).

31. Zañudo, J. G. T. & Albert, R. An effective network reduction approach to find the dynamical repertoire of discrete dynamic networks. Chaos: An Interdisciplinary Journal of Nonlinear Science 23, 025111 (2013).

32. Gan, X. & Albert, R. General method to find the attractors of discrete dynamic models of biological systems. Physical Review E 97, (2018).

33. Ravasz, E., Somera, A. L., Mongru, D. A., Oltvai, Z. N. & Barabási, A. L. Hierarchical organization of modularity in metabolic networks. Science 297, 1551–1555 (2002).

34. Ihmels, J. et al. Revealing modular organization in the yeast transcriptional network. Nat. Genet. 31, 370–377 (2002).

35. Ma, H.-W., Zhao, X.-M., Yuan, Y.-J. & Zeng, A.-P. Decomposition of metabolic network into functional modules based on the global connectivity structure of reaction graph. Bioinformatics 20, 1870–1876 (2004).

36. Papin, J. A., Reed, J. L. & Palsson, B. O. Hierarchical thinking in network biology: the unbiased modularization of biochemical networks. Trends Biochem. Sci. 29, 641–647 (2004).

37. Tanay, A., Sharan, R., Kupiec, M. & Shamir, R. Revealing modularity and organization in the yeast molecular network by integrated analysis of highly heterogeneous genomewide data. Proc. Natl. Acad. Sci. U. S. A. 101, 2981–2986 (2004).

38. Neumann, B. et al. Phenotypic profiling of the human genome by time-lapse microscopy reveals cell division genes. Nature 464, 721–727 (2010).

39. Vitorino, P. & Meyer, T. Modular control of endothelial sheet migration. Genes Dev. 22, 3268–3281 (2008).

40. Wynn, M. L., Consul, N., Merajver, S. D. & Schnell, S. Logic-based models in systems biology: a predictive and parameter-free network analysis method. Integrative Biology 4, 1323 (2012).

41. Abou-Jaoudé, W. et al. Logical Modeling and Dynamical Analysis of Cellular Networks. Frontiers in Genetics 7, (2016).

42. Bloomingdale, P., Nguyen, V. A., Niu, J. & Mager, D. E. Boolean network modeling in systems pharmacology. Journal of Pharmacokinetics and Pharmacodynamics 45, 159–180 (2018).

43. Zañudo, J. G. T., Steinway, S. N. & Albert, R. Discrete dynamic network modeling of oncogenic signaling: Mechanistic insights for personalized treatment of cancer. Current Opinion in Systems Biology 9, 1–10 (2018).

44. Rozum, J. C. & Albert, R. Identifying (un)controllable dynamical behavior in complex networks. PLOS Computational Biology 14, e1006630 (2018).

45. Wang, R.-S. & Albert, R. Elementary signaling modes predict the essentiality of signal transduction network components. BMC Systems Biology 5, (2011).

46. Novak, B., Tyson, J. J., Gyorffy, B. & Csikasz-Nagy, A. Irreversible cell-cycle transitions are due to systems-level feedback. Nature Cell Biology 9, 724–728 (2007).

47. Burdon, T., Smith, A. & Savatier, P. Signalling, cell cycle and pluripotency in embryonic stem cells. Trends Cell Biol. 12, 432–438 (2002).

48. Marques-Pita, M. & Rocha, L. M. Canalization and control in automata networks: body segmentation in Drosophila melanogaster. PLoS One 8, e55946 (2013).

49. Li, Y., Adeyeye, J. O., Murrugarra, D., Aguilar, B. & Laubenbacher, R. Boolean nested canalizing functions: A comprehensive analysis. Theoretical Computer Science 481, 24–36 (2013).

50. Freeman, L. C. A Set of Measures of Centrality Based on Betweenness. Sociometry. 40, 35 (1977).

51. Sha, W. et al. Hysteresis drives cell-cycle transitions in Xenopus laevis egg extracts. Proceedings of the National Academy of Sciences 100, 975–980 (2003).

52. Pomerening, J. R., Sontag, E. D. & Ferrell, J. E. Building a cell cycle oscillator: hysteresis and bistability in the activation of Cdc2. Nature Cell Biology 5, 346–351 (2003).

53. Novak, B. & Tyson, J. J. Numerical analysis of a comprehensive model of M-phase control in Xenopus oocyte extracts and intact embryos. J. Cell Sci. 106 (Pt 4), 1153–1168 (1993).

54. He, E. et al. System-level feedbacks make the anaphase switch irreversible. Proceedings of the National Academy of Sciences 108, 10016–10021 (2011).

55. Kapuy, O., He, E., Uhlmann, F. & Novák, B. Mitotic exit in mammalian cells. Mol. Syst. Biol. 5, 324 (2009).

56. Potapova, T. A., Daum, J. R., Byrd, K. S. & Gorbsky, G. J. Fine tuning the cell cycle: activation of the Cdk1 inhibitory phosphorylation pathway during mitotic exit. Mol. Biol. Cell 20, 1737–1748 (2009).

57. PageRank Algorithm, 1998; Brin, Page. SpringerReference doi:10.1007/springerreference_57796

58. Yang, G., Zañudo, J. G. T. & Albert, R. Target Control in Logical Models Using the Domain of Influence of Nodes. Frontiers in Physiology 9, (2018).

