## Supplemental Texts for "A feedback loop of conditionally stable circuits drives the cell cycle from checkpoint to checkpoint"

**Supplementary Text S1. Regulatory functions of the Phase Switch module and the Phase Switch Oscillator**

**Regulatory functions of the Phase Switch module**

f_Cdc25A_=CyclinA and not Cdh1

f_Mad2_= (not pAPC and CyclinB and Cdk1) or (not Cdc20 and CyclinB and Cdk1)

f_Wee1_= (not CyclinA and not CyclinB) or not Cdk1

f_Cdc25C_= CyclinA or (CyclinB and Cdk1)

f_Cdh1_= (not CyclinA and not CyclinB) or (not CyclinA and not Cdk1)

f_Cdk1_= (Cdc25C and CyclinA and Cdk1) or (Cdc25C and CyclinB and Cdk1) or (Cdc25C and CyclinA and not Wee1) or (Cdc25C and CyclinB and not Wee1)

f_CyclinA_= (not pAPC and not Cdh1 and Cdc25A) or (not pAPC and not UbcH10 and Cdc25A) or (not pAPC and not Cdh1 and CyclinA) or (not pAPC and not Cdh1 and CyclinA)

f_CyclinB_= (not pAPC and not Cdh1) or (not Cdc20 and not Cdh1)

f_pAPC_= (pAPC and Cdc20) or (CyclinB and Cdk1)

f_Cdc20_= pAPC and not Cdh1 and not Mad2

f_UbcH10_= not Cdh1 or (UbcH10 and Cdc20) or (UbcH10 and CyclinA) or (UbcH10 and CyclinB)

**Regulatory functions of the nodes of the Phase Switch module in the full cell cycle model**

Regulators that are outside of the Phase Switch module are indicated in bold.

f_Cdc25A_= (**E2F1** and **CyclinE**) or (**E2F1** and CyclinA) or (CyclinA and not Cdh1 and **CyclinE**)

f_Mad2_= (**4N_DNA** and not pAPC and CyclinB and Cdk1 and not **Metaphase**) or (**4N_DNA** and not Cdc20 and CyclinB and Cdk1 and not **Metaphase**)

f_Wee1_= (not CyclinA and not CyclinB and **Replication**) or (not Cdk1 and **Replication**)

f_Cdc25C_= CyclinA or (CyclinB and Cdk1)

f_Cdh1_= (not CyclinA and not CyclinB) or (not CyclinA and not Cdk1)

f_Cdk1_= (Cdc25C and CyclinA and Cdk1) or (Cdc25C and CyclinB and Cdk1) or (Cdc25C and CyclinA and not Wee1) or (Cdc25C and CyclinB and not Wee1)

f_CyclinA_= (not pAPC and not Cdh1 and Cdc25A and **E2F1**) or (not pAPC and not UbcH10 and Cdc25A and **E2F1**) or (not pAPC and not Cdh1 and CyclinA) or (not pAPC and not Cdh1 and CyclinA)

f_CyclinB_= (not pAPC and not Cdh1) or (not Cdc20 and not Cdh1)

f_pAPC_= (pAPC and Cdc20) or (CyclinB and Cdk1)

f_Cdc20_= pAPC and not Cdh1 and not Mad2

f_UbcH10_= not Cdh1 or (UbcH10 and Cdc20) or (UbcH10 and CyclinA) or (UbcH10 and CyclinB)

**Regulatory functions of the Phase Switch Oscillator (after substituting Cdc25A=1, Mad2=Wee1=0)**

f_Cdc25C_= CyclinA or (CyclinB and Cdk1)

f_Cdh1_= (not CyclinA and not CyclinB) or (not CyclinA and not Cdk1)

f_Cdk1_= (Cdc25C and CyclinA) or (Cdc25C and CyclinB)

f_CyclinA_= (not pAPC and not Cdh1) or (not pAPC and not UbcH10)

f_CyclinB_= (not pAPC and not Cdh1) or (not Cdc20 and not Cdh1)

f_pAPC_= (pAPC and Cdc20) or (CyclinB and Cdk1)

f_Cdc20_= pAPC and not Cdh1

f_UbcH10_= not Cdh1 or (UbcH10 and Cdc20) or (UbcH10 and CyclinA) or (UbcH10 and CyclinB)

**Regulatory functions of the Phase Switch Oscillator embodied in the expanded network**

f_Cdc25C_= CyclinA or (CyclinB and Cdk1)

f_~Cdc25C_= (~CyclinA and ~CyclinB) or (~CyclinA and Cdk1)

f_Cdh1_= (~CyclinA and ~CyclinB) or (~CyclinA and ~Cdk1)

f_~Cdh1_= CyclinA or (CyclinB and Cdk1)

f_Cdk1_= (Cdc25C and CyclinA) or (Cdc25C and CyclinB)

f_~Cdk1_= ~Cdc25C or (~CyclinA and ~CyclinB)

f_CyclinA_= (~pAPC and ~Cdh1) or (~pAPC and ~UbcH10)

f_~CyclinA_= pAPC or (Cdh1 and UbcH10)

f_CyclinB_= (~pAPC and ~Cdh1) or (~Cdc20 and ~Cdh1)

f_~CyclinB_= (pAPC and Cdc20) or Cdh1

f_pAPC_= (pAPC and Cdc20) or (CyclinB and Cdk1)

f_~pAPC_= (~pAPC and ~CyclinB) or (~Cdc20 and ~CyclinB) or (~pAPC and ~Cdk1) or (~Cdc20 and ~Cdk1)

f_Cdc20_= pAPC and ~Cdh1

f_~Cdc20_= ~pAPC or Cdh1

f_UbcH10_= ~Cdh1 or (UbcH10 and Cdc20) or (UbcH10 and CyclinA) or (UbcH10 and CyclinB)

f_~UbcH10_= (Cdh1 and ~UbcH10) or (Cdh1 and ~Cdc20 and ~CyclinA and ~CyclinB)

**Supplementary Text S2. Detailed description of the complex attractor of the Phase**

**Switch Oscillator**

In the following we describe the state transitions involved in the complex attractor of the PSO, starting with the state closest to the G0/G1 attractor. As the G0/G1 state overlaps the G2 state in three node states (see Supplementary Table S5), five nodes need to change from the close-to-G0/G1 to the close-to-G2 state: CyclinB, UbcH10, CyclinA, Cdc25C need to turn on, Cdh1 needs to turn off. CyclinA turns on first (in a transition shared with synchronous update, see Figure 4). In most trajectories this leads to Cdh1 turning off and Cdc25C turning on, in either order. Next, CyclinB and UbcH10 turn on in either order, with the Cdk1-on transition mixed in. A considerable fraction of the asynchronous trajectories mixes the state changes of Cdh1, Cdc25C, CyclinB, UbcH10, Cdk1, and thus form a transition between the state with overlap trio (7,4,1) and the post-G2 state that skips the state (5,6,3). There also is a small fraction of trajectories wherein the state change of UbcH10 is delayed; these trajectories skip the post-G2 state. The rest of the trajectories follow the two steps of the synchronous update.   The state closest to the G2 attractor is contained in half of the trajectories between the state (7,4,1) and the post-G2 state.

As the state closest to G2 overlaps the state closest to SAC in five node states, only three nodes need to change state to switch from the G2 to SAC state: pAPC and Cdk1 need to turn on, while CyclinA needs to turn off. Cdk1 turns on first, then pAPC (in a transition shared by synchronous update). Then CyclinA turns off and Cdc20 turns on, in either order, marking a transition between the near-SAC state to the post-SAC state. The SAC state is visited by the trajectories wherein CyclinA turns off first and then Cdc20 turns on (approximately half of the total trajectories, see Supplementary Table S5). There are also a few trajectories where the turning on of UbcH10 is delayed, and follows the turning on of pAPC, Cdc20, or the turning off of CyclinA.

Finally, six nodes need to change state from the SAC to the G0/G1 state: pAPC, Cdk1, CyclinB, UbcH10, and Cdc25C need to turn off, while Cdh1 needs to turn on. In addition, Cdc20 first turns on (during the transition from the near-SAC to the post-SAC state), then it turns off during the approach to the G0/G1 state.  CyclinB turns off in a transition shared by the synchronous cycle, between the post-SAC state and the state with attractor overlap (2,3,6). It is also possible that the turning off of Cyclin B precedes that of CyclinA, creating trajectories from the near-SAC state to (2,3,6) that skip the post-SAC state. Following the state (2,3,6), the Cdh1-on, Cdc25C-off and Cdk1-off transitions can occur in variable order. Cdc20, pAPC, and UbcH10 turn off in variable order following the activation of Cdh1, in trajectories that reach the G0/G1 atate with or without visiting the states (5,2,3) and (6,3,4).

**Supplementary Text S3. Detailed description of the agreement of the expanded network and complex attractor of the Phase Switch Oscillator**

To illustrate the correspondence between the expanded network (Supplementary Figure S4) and the complex attractor (Figure 4) of the Phase Switch Oscillator, we go around the state transition backbone, starting with the G0/G1 state.  The expanded network indicates that the condition for CyclinA to turn on (i.e. the condition for the virtual node CyclinA to be reached) is that UbcH10 and pAPC are simultaneously off (this is expressed by a composite node whose regulators are ~UbcH10 and ~pAPC) or Cdc20 and Cdh1 are simultaneously off. The first of these conditions is satisfied in the G0/G1 state, thus CyclinA will turn on with certainty, in agreement with and explaining the probability 1 of the edge between the G0/G1 state and the state with overlap trio (7,4,1). Cdc25C turns on and Cdh1 turns off in arbitrary order between the state with overlap (7,4,1) and the state (5,6,3) because Cdc25C and ~Cdh1 are driven by CyclinA (i.e. there is an edge from CyclinA to Cdc25C and an edge from CyclinA to ~Cdh1). UbcH10 is driven by ~Cdh1, thus its state transition follows that of Cdh1. Cdk1 turns on if CyclinA and Cdc25C are simultaneously on, thus its state transition follows the turning on of Cdc25C. The turning on of CyclinB needs the simultaneous OFF state of Cdh1 and pAPC or of Cdh1 and Cdc20 (indicated by two separate composite nodes). Both pAPC and Cdc20 are off in the state (7,4,1), thus the state change of CyclinB follows that of Cdh1. The multiple orders in which the previously described five nodes can change state induces the possibility of paths from the state (7,4,1) to  the post-G2 state that skip the state (5,6,3).

The turning on of pAPC necessitates CyclinB and Cdk1 (see composite node), both of which are present in the post-G2 state, thus this will happen with high probability. The expanded network’s subgraph that starts with ~pAPC and CyclinA, contains Cdc25C, ~Cdh1, Cdk1, CyclinB as well as 3 composite nodes, and converges on pAPC, expresses the minimal logical condition for the mediated self-inhibition of pAPC. In other words, the logic domain of influence (LDOI) of the node set {~pAPC, CyclinA} contains pAPC. The turning off of CyclinA is driven by pAPC, and the turning on of Cdc20 is driven by the combination of pAPC and ~Cdh1. Cdc20 contributes to the self-sustained expression of pAPC and UbcH10 by being the condition of their conditionally stable motifs. The subgraph of the expanded network that starts with CyclinA and CyclinB and contains Cdc25C, Cdk1, pAPC and three composite nodes, ends in ~CyclinA. Thus, the LDOI of {CyclinA, CyclinB} contains ~CyclinA. CyclinB turns off if Cdc20 and pAPC are simultaneously present.

The subgraph that starts with CyclinB and Cdk1 and contains ~Cdh1, pAPC (both of which are regulated by the same composite node), Cdc20 and two additional composite nodes, and ends in ~CyclinB, expresses the mediated self-inhibition of CyclinB. Thus, the LDOI of {CyclinB, Cdk1} contains ~CyclinB.  This subgraph overlaps the C11 conditionally stable motif, which includes the virtual nodes ~Cdh1, CyclinB, Cdk1, Cdc25C and is conditioned either on ~pAPC or ~Cdc20. We can see that this CSM drives its own destabilization, as the composite node that maintains ~Cdh1 also drives pAPC, which negates one of the conditions. pAPC together with ~Cdh1 drives Cdc20, which negates the other condition.

The virtual nodes ~CyclinA and ~CyclinB together regulate a composite node that drives all three of Cdh1, ~Cdk1 and ~Cdc25C. This is the reason these three nodes switch states in arbitrary order between the state marked (2,3,6) and (5,2,3), while with synchronous update the second state is a direct successor of the first.  Cdh1 is sufficient to turn off Cdc20; this in certain trajectories can happen before the turning off of Cdc25C or Cdk1, contributing to the possibility of a trajectory from state (2,3,6) directly to (6,3,4). The off state of Cdc20, combined with ~CyclinB or ~Cdk1, drive ~pAPC. The turning off of UbcH10 necessitates the combination of ~CyclinA, ~CyclinB, Cdh1, ~Cdc20. Since all of these node states were reached in the previous steps, the turning off of pAPC and UbcH10 can happen in arbitrary order. The off state of UbcH10 can be sustained with the help of Cdh1 (bidirectional edge in the expanded network). The simultaneous off state of UbcH10 and pAPC is sufficient to turn CyclinA on. Thus, the subgraph that starts with ~CyclinA and ~CyclinB, contains Cdh1, ~Cdk1, ~Cdc20, ~UbcH10, ~pAPC and four composite nodes, and ends in CyclinA, expresses the mediated contradiction in CyclinA. In other words, the LDOI of {~CyclinA, ~CyclinB} contains CyclinA.
