## Supplemental Tables for "A feedback loop of conditionally stable circuits drives the cell cycle from checkpoint to checkpoint"

**Supplementary Table S1. The three fixed point attractors of the Phase Switch**, using the same notations as Deritei et al 2016. **G0/G1** represents the quiescent or first growth phase of the cell before the Restriction Point. Both in the attractor state and the mammalian cell all molecules are inactive, except for Cdh1. The difference between the global G0 state of the cell cycle model and the G0/G1 state of the Phase Switch is the ON state of Wee1, which is not expressed in cells in G0/G1. **G2** represents the second growth phase and the locked-in state of the cell before the DNA damage checkpoint is cleared. The main driver molecules of the cell cycle are on in this phase, such as CyclinA and CyclinB. The activation of Wee1 is critical here because its mutual inhibition with Cdk1 blocks the system from passing the DNA damage checkpoint. The one biological inaccuracy is the active state of Cdc25C, which is primed by CyclinA/Cdk2 activity (represented by the CyclinA node), but is not fully active in the presence of Wee1^S1^. The model’s logic compensates for this by requiring both the presence of Cdc25C and the absence of Wee1 to allow Cdk1 activation. **SAC** corresponds to the phase where the spindle that will pull sister chromatids apart is assembled, thus as its name suggests represents the point before passing the spindle assembly checkpoint. Here the early driver of the cycle, CyclinA, is already turned off, while the CyclinB/Cdk1 complexes are still active. The checkpoint is enforced by the activity of Mad2, which remains on as long as there is even a single unattached kinetochore on the cell’s replicated chromosomes. This checkpoint guards cells from erroneous chromosome segregation into the two daughter cells.

|  | **Cdc25C** | **CyclinA** | **Cdk1** | **CyclinB** | **Cdh1** | **pAPC** | **Cdc20** | **UbcH10** | **Mad2** | **Wee1** | **Cdc25A** |
| --- | --- | --- | --- | --- | --- | --- | --- | --- | --- | --- | --- |
| **G0/G1** | 0 | 0 | 0 | 0 | 1 | 0 | 0 | 0 | 0 | 1 | 0 |
| **G2** | 1 | 1 | 0 | 1 | 0 | 0 | 0 | 1 | 0 | 1 | 1 |
| **SAC** | 1 | 0 | 1 | 1 | 0 | 1 | 0 | 1 | 1 | 0 | 0 |

**Supplementary Table S2. Probability of state transitions among pairs of states of the synchronous limit cycle in the complex attractor found by general asynchronous update.** The first two columns indicate the start and end state. The paths of the complex attractor (indicated in the fourth column) are grouped by the nodes that change state along the path. The turning on of a node is indicated by the node name and the turning off is indicated by preceding the node name by ~. The third column separately indicates the probability of the path that involves the same node state changes as the synchronous limit cycle and the probability of longer paths that have the same end state. The latter probability is a small correction to the former.

| **Starting state** | **End state** | **Transition probability** | **States that change on the path (ignoring permutations)** |
| --- | --- | --- | --- |
| (8, 3, 2) | (7, 4, 1) | 1 | CyclinA |
| (1, 4, 7) | (2, 3, 6) | 1 | ~CyclinB |
| (5, 2, 3) | (6, 3, 4) | 1 | ~Cdc20 |
| (6, 3, 4) | (8, 3, 2) | 1 | ~pAPC, ~UbcH10 |
| (2, 7, 6) | (1, 6, 7) | 1 | pAPC |
| (5, 6, 3) | (2, 7, 6) | 0.8333 | Cdk1, CyclinB, UbcH10 |
|  |  | 0.0003 | CyclinB, Cdk1, pAPC, ~CyclinA, Cdc20, ~CyclinB, Cdh1, ~Cdc20, ~pAPC, CyclinA, ~Cdh1, UbcH10, CyclinB |
|  |  |  | CyclinB, Cdk1, pAPC, ~CyclinA, Cdc20, ~CyclinB, Cdh1, ~Cdc20, ~Cdk1, ~pAPC, CyclinA, Cdk1, ~Cdh1, UbcH10, CyclinB |
|  |  |  | Cdk1, CyclinB, pAPC, ~CyclinA, Cdc20, ~CyclinB, Cdh1, ~Cdc20, ~pAPC, ~Cdc25C, CyclinA, ~Cdh1, UbcH10, CyclinB, Cdc25C |
| (1, 6, 7) | (1, 4, 7) | 0.75 | ~CyclinA, Cdc20 |
| (2, 3, 6) | (5, 2, 3) | 0.6111 | ~Cdc25C, Cdh1, ~Cdk1 |
|  |  | 0.0002 | ~Cdc25C, Cdh1, ~Cdc20, ~pAPC, ~UbcH10, CyclinA, ~Cdh1, UbcH10, CyclinB, pAPC, ~CyclinA, Cdc20, ~Cdk1, ~CyclinB, Cdh1 |
| (7, 4, 1) | (2, 7, 6) | 0.5000 | ~Cdh1, CyclinB, UbcH10, Cdc25C, Cdk1 |
|  |  | 10^-5^ | ~Cdh1, CyclinB, Cdc25C, Cdk1, pAPC, ~CyclinA, Cdc20, ~CyclinB, Cdh1, ~Cdc20, ~pAPC, CyclinA, ~Cdh1, UbcH10, CyclinB |
| (7, 4, 1) | (5, 6, 3) | 0.4166 | ~Cdh1, Cdc25C |
|  |  | 0.0001 | ~Cdh1, CyclinB, Cdc25C, Cdk1, pAPC, ~CyclinA, Cdc20, ~CyclinB, Cdh1, ~Cdc20, ~Cdk1, ~pAPC, CyclinA, ~Cdh1 |
|  |  |  | Cdc25C, Cdk1, ~Cdh1, CyclinB, pAPC, ~CyclinA, Cdc20, ~CyclinB, Cdh1, ~Cdc20, ~pAPC, ~Cdc25C, CyclinA, ~Cdh1, ~Cdk1, Cdc25C |
|  |  |  | ~Cdh1, CyclinB, Cdc25C, Cdk1, pAPC, ~CyclinA, Cdc20, ~CyclinB, ~Cdk1, UbcH10, Cdh1, ~Cdc20, ~UbcH10, ~pAPC, CyclinA, ~Cdh1 |
| (1, 6, 7) | (2, 3, 6) | 0.25 | Cdc20, ~CyclinB, ~CyclinA |
| (2, 3, 6) | (8, 3, 2) | 0.2006 | ~Cdc25C, Cdh1, ~Cdc20, ~pAPC, ~UbcH10, ~Cdk1 |
|  |  | 0.0002 | ~Cdc25C, Cdh1, ~Cdc20, ~pAPC, ~UbcH10, CyclinA, ~Cdh1, CyclinB, pAPC, ~CyclinA, ~Cdk1, Cdh1, ~pAPC, ~CyclinB |
|  |  |  | ~Cdc25C, Cdh1, ~Cdc20, ~pAPC, ~UbcH10, CyclinA, ~Cdh1, CyclinB, pAPC, ~CyclinA, Cdc20, ~Cdk1, ~CyclinB, Cdh1, ~Cdc20, ~pAPC |
|  |  |  | ~Cdc25C, Cdh1, ~Cdc20, ~pAPC, ~UbcH10, CyclinA, ~Cdh1, CyclinB, pAPC, ~Cdk1, Cdc25C, ~CyclinA, ~pAPC, Cdh1, ~CyclinB, ~Cdc25CX |
|  |  |  | ~Cdc25C, Cdh1, ~Cdc20, ~pAPC, ~UbcH10, CyclinA, ~Cdh1, UbcH10, CyclinB, pAPC, ~CyclinA, ~Cdk1, ~pAPC, Cdh1, ~CyclinB, ~UbcH10 |
| (2, 3, 6) | (6, 3, 4) | 0.1111 | ~Cdc25C, Cdh1, ~Cdc20, ~Cdk1 |
|  |  | 10^-5^ | ~Cdc25C, Cdh1, ~Cdc20, ~pAPC, ~UbcH10, CyclinA, ~Cdh1, UbcH10, CyclinB, pAPC, ~CyclinA, ~Cdk1, Cdh1, ~CyclinB |
|  |  |  | ~Cdc25C, Cdh1, ~Cdc20, ~pAPC, ~UbcH10, CyclinA, ~Cdh1, UbcH10, CyclinB, pAPC, ~CyclinA, Cdc20, ~Cdk1, Cdh1, ~Cdc20, ~CyclinB |
|  |  |  | ~Cdc25C, Cdh1, ~Cdc20, ~pAPC, ~UbcH10, CyclinA, ~Cdh1, UbcH10, CyclinB, pAPC, ~Cdk1, Cdc25C, ~CyclinA, Cdh1, ~CyclinB, ~Cdc25C |
| (5, 6, 3) | (1, 4, 7) | 0.0601 | Cdk1, CyclinB, pAPC, ~CyclinA, UbcH10, Cdc20 |
| (5, 6, 3) | (1, 6, 7) | 0.0555 | Cdk1, CyclinB, pAPC, UbcH10 |
|  |  | 10^-5^ | Cdk1, CyclinB, pAPC, ~CyclinA, Cdc20, ~CyclinB, Cdh1, ~Cdc20, ~pAPC, ~Cdc25C, CyclinA, ~Cdh1, UbcH10, CyclinB, pAPC, Cdc25C |
| (2, 3, 6) | (2, 7, 6) | 0.0351 | ~Cdc25C, Cdh1, ~Cdc20, ~pAPC, ~UbcH10, CyclinA, ~Cdh1, UbcH10, CyclinB, Cdc25C |
|  |  |  | ~Cdc25C, Cdh1, ~Cdc20, ~pAPC, ~UbcH10, CyclinA, ~Cdh1, UbcH10, CyclinB, ~Cdk1, Cdc25C, Cdk1 |
|  |  |  | ~Cdc25C, Cdh1, ~Cdc20, ~pAPC, ~UbcH10, CyclinA, ~Cdh1, UbcH10, CyclinB, pAPC, ~Cdk1, ~pAPC, Cdc25C, Cdk1 |
|  |  |  | ~Cdc25C, Cdh1, ~Cdc20, ~pAPC, ~UbcH10, CyclinA, ~Cdh1, UbcH10, CyclinB, pAPC, ~CyclinA, ~Cdk1, ~pAPC, CyclinA, Cdc25C, Cdk1 |
|  |  |  | Cdh1, ~Cdk1, ~Cdc20, ~UbcH10, ~pAPC, CyclinA, Cdk1, ~Cdh1, CyclinB, UbcH10XCdh1, ~Cdc20, ~UbcH10, ~pAPC, CyclinA, ~Cdh1, CyclinB, UbcH10 |
| (7, 4, 1) | (1, 4, 7) | 0.0300 | ~Cdh1, CyclinB, Cdc25C, Cdk1, pAPC, ~CyclinA, UbcH10, Cdc20 |
| (7, 4, 1) | (1, 6, 7) | 0.0277 | ~Cdh1, CyclinB, Cdc25C, Cdk1, pAPC, UbcH10 |
| (5, 6, 3) | (2, 3, 6) | 0.0266 | Cdk1, CyclinB, pAPC, ~CyclinA, Cdc20, ~CyclinB, UbcH10 |
| (2, 3, 6) | (5, 6, 3) | 0.0187 | Cdh1, ~Cdk1, ~Cdc20, ~UbcH10, ~pAPC, CyclinA, ~Cdh1 |
|  |  |  | ~Cdc25C, Cdh1, ~Cdc20, ~pAPC, ~UbcH10, CyclinA, ~Cdh1, ~Cdk1, Cdc25C |
| (5, 6, 3) | (8, 3, 2) | 0.0138 | Cdk1, CyclinB, pAPC, ~CyclinA, Cdc20, ~CyclinB, Cdh1, ~Cdc20, ~Cdk1, ~pAPC, ~Cdc25C |
|  |  | 0.0006 | Cdk1, CyclinB, pAPC, ~CyclinA, Cdc20, ~CyclinB, ~Cdc25C, UbcH10, Cdh1, ~Cdc20, ~pAPC, ~UbcH10, ~Cdk1 |
| (7, 4, 1) | (2, 3, 6) | 0.0133 | ~Cdh1, CyclinB, Cdc25C, Cdk1, pAPC, ~CyclinA, Cdc20, ~CyclinB, UbcH10 |
| (2, 3, 6) | (7, 4, 1) | 0.0118 | ~Cdc25C, Cdh1, ~Cdc20, ~pAPC, ~UbcH10, CyclinA, ~Cdk1 |
|  |  |  | ~Cdc25C, Cdh1, ~Cdc20, ~pAPC, ~UbcH10, CyclinA, ~Cdh1, CyclinB, pAPC, ~CyclinA, ~Cdk1, Cdh1, ~pAPC, CyclinA, ~CyclinB |
| (7, 4, 1) | (8, 3, 2) | 0.0072 | ~Cdh1, CyclinB, Cdc25C, Cdk1, pAPC, ~CyclinA, Cdc20, ~CyclinB, Cdh1, ~Cdc20, ~Cdk1, ~pAPC, ~Cdc25C |
|  |  |  | ~Cdh1, CyclinB, Cdc25C, Cdk1, pAPC, ~CyclinA, Cdc20, ~CyclinB, ~Cdc25C, UbcH10, Cdh1, ~Cdc20, ~pAPC, ~UbcH10, ~Cdk1 |
| (5, 6, 3) | (5, 2, 3) | 0.0067 | Cdk1, CyclinB, pAPC, ~CyclinA, Cdc20, ~CyclinB, ~Cdc25C, UbcH10, Cdh1, ~Cdk1 |
| (2, 3, 6) | (1, 4, 7) | 0.0040 | Cdh1, ~Cdc20, ~UbcH10, ~pAPC, CyclinA, ~Cdh1, CyclinB, pAPC, ~CyclinA, UbcH10, Cdc20 |
|  |  |  | Cdh1, ~Cdk1, ~Cdc20, ~UbcH10, ~pAPC, CyclinA, Cdk1, ~Cdh1, CyclinB, pAPC, ~CyclinA, UbcH10, Cdc20 |
|  |  |  | ~Cdc25C, Cdh1, ~Cdc20, ~pAPC, ~UbcH10, CyclinA, ~Cdh1, UbcH10, CyclinB, pAPC, ~CyclinA, Cdc25C, Cdc20 |
|  |  |  | ~Cdc25C, Cdh1, ~Cdc20, ~pAPC, ~UbcH10, CyclinA, ~Cdh1, UbcH10, CyclinB, pAPC, ~Cdk1, Cdc20, Cdc25C, Cdk1, ~CyclinA |
| (2, 3, 6) | (1, 6, 7) | 0.0036 | Cdh1, ~Cdc20, ~UbcH10, ~pAPC, CyclinA, ~Cdh1, CyclinB, pAPC, UbcH10 |
|  |  |  | ~Cdc25C, Cdh1, ~Cdc20, ~pAPC, ~UbcH10, CyclinA, ~Cdh1, UbcH10, CyclinB, pAPC, Cdc25C |
|  |  |  | Cdh1, ~Cdk1, ~Cdc20, ~UbcH10, ~pAPC, CyclinA, Cdk1, ~Cdh1, CyclinB, pAPC, UbcH10 |
|  |  |  | ~Cdc25C, Cdh1, ~Cdc20, ~pAPC, ~UbcH10, CyclinA, ~Cdh1, UbcH10, CyclinB, pAPC, ~Cdk1, Cdc25C, Cdk1 |
|  |  |  | ~Cdc25C, Cdh1, ~Cdc20, ~pAPC, ~UbcH10, CyclinA, ~Cdh1, CyclinB, pAPC, ~Cdk1, Cdc25C, ~pAPC, Cdk1, pAPC, UbcH10 |
| (7, 4, 1) | (5, 2, 3) | 0.0034 | ~Cdh1, CyclinB, Cdc25C, Cdk1, pAPC, ~CyclinA, Cdc20, ~CyclinB, ~Cdc25C, UbcH10, Cdh1, ~Cdk1 |
| (5, 6, 3) | (6, 3, 4) | 0.0005 | Cdk1, CyclinB, pAPC, ~CyclinA, Cdc20, ~CyclinB, ~Cdc25C, UbcH10, Cdh1, ~Cdc20, ~Cdk1 |
| (5, 6, 3) | (7, 4, 1) | 0.0004 | Cdk1, CyclinB, pAPC, ~CyclinA, Cdc20, ~CyclinB, Cdh1, ~Cdc20, ~pAPC, ~Cdc25C, CyclinA, ~Cdk1 |
|  |  |  | Cdk1, CyclinB, pAPC, ~CyclinA, Cdc20, ~CyclinB, ~Cdc25C, UbcH10, Cdh1, ~Cdc20, ~pAPC, ~UbcH10, CyclinA, ~Cdk1 |
| (7, 4, 1) | (6, 3, 4) | 0.0002 | ~Cdh1, CyclinB, Cdc25C, Cdk1, pAPC, ~CyclinA, Cdc20, ~CyclinB, ~Cdc25C, UbcH10, Cdh1, ~Cdc20, ~Cdk1 |

**Supplementary Table S3. The conditionally stable motifs (CSMs) of the PSO network that cause their own destabilization**. The first column is the identifier of the CSM. The second column contains the node states (virtual nodes on the expanded network where ~ refers to the 0 state) making up the CSM. The third column is the state that serves as the condition that needs to be sustained for the CSM to be stable. The opposite state is contained in the logic domain of influence of the CSM. The fourth column indicates the cases where the CSM corresponds to a motif of the Phase Switch (shown on Figure 2). Such destabilization does not happen in the Phase Switch because the stable motifs either stabilize, or violate, the condition of each CSM (see Figure 2).

| **Name** | **Virtual nodes of the conditionally stable motif** | **Virtual node that serves as condition** | **Corresponding Phase Switch motif** |
| --- | --- | --- | --- |
| C0 | ~Cdk1, ~Cdc5C | ~CyclinA | P3 |
| C1 | ~CyclinB, Cdh1 | ~CyclinA | P4 |
| C2 | ~CyclinB, Cdh1, ~Cdk1 | ~CyclinA |  |
| C3 | ~CyclinA, Cdh1, ~CyclinB | UbcH10 |  |
| C4 | ~CyclinA, ~Cdk1, Cdh1, ~CyclinB | UbcH10 |  |
| C5 | ~CyclinA, ~Cdk1, Cdh1, ~Cdc25C | UbcH10 |  |
| C8 | ~UbcH10 | Cdh1 |  |
| C9 | ~Cdh1, CyclinA | ~pAPC | P6 |
| C10 | Cdk1, Cdc25C | CyclinB |  |
| C11 | ~Cdh1, CyclinB, Cdk1, Cdc25C | ~Cdc20 or ~pAPC | P2’ |
| C13 | pAPC | Cdc20 |  |
| C14 | pAPC, Cdc20 | ~Cdh1 |  |

**Supplementary Table S4. List of attractors that arise when the node state listed in the “Intervention” column is held fixed.** Columns denoted by node names give the steady state value of the respective node (grey background highlights locked nodes), and the Overlap column gives the overlap of this steady state with each of the three Phase Switch attractors in the order (G0/G1, G2, SAC). The next column indicates the closest Phase Switch attractor(s). Arrows to / from phenotypes in parentheses point to other attractors with near-maximum overlap, indicating that the network is stuck close to the boundary between two Phase Switch states. The final column gives the conditionally stable motifs for which the intervention satisfied a condition; these become stable motifs in the modified system and underlie the resulting attractor. A node state listed in the final column indicates that the intervention is a sufficient condition for setting this state. The interventions UbcH10=0, Cdc25c=1, and Cdk1=1 each result in a single complex attractor in which each remaining node oscillates; in this case no conditionally stable motif becomes a stable motif, and the steady state overlap is not well-defined.

| **Intervention** | **Attractors of the altered system** | | | | | | | | | **Overlap (G0/G1, G2, SAC)** | **Closest Phase Switch Attractor(s)** | **CSMs stabilized by the intervention** |
| --- | --- | --- | --- | --- | --- | --- | --- | --- | --- | --- | --- | --- |
|  |  | **CyclinA** | **CyclinB** | **Cdc20** | **Cdc25c** | **Cdh1** | **Cdk1** | **pAPC** | **UbcH10** |  |  |  |
| UbcH10=0 | Oscillating | Osc. | Osc. | Osc. | Osc. | Osc. | Osc. | Osc. | 0 | N/A |  | None |
| Cdc25c=1 |  | Osc. | Osc. | Osc. | 1 | Osc. | Osc. | Osc. | Osc. | N/A |  | None |
| Cdk1=1 |  | Osc. | Osc. | Osc. | Osc. | Osc. | 1 | Osc. | Osc. | N/A |  | None |
| CyclinA=0 | Monostable | 0 | 0 | 0 | 0 | 1 | 0 | 0 | 0 | (8,3,2) | G0/G1 | C0/P3, C1/P4, C2 |
| pAPC=0 |  | 0 | 0 | 0 | 0 | 1 | 0 | 0 | 0 | (7,2,3) | G0/G1 | C9/P6, Cdc20=0 |
| UbcH10=1 |  | 0 | 0 | 0 | 0 | 1 | 0 | 0 | 1 | (7,4,3) | G0/G1 | C3, C4, C5 |
| Cdh1=1 |  | 1 | 0 | 0 | 1 | 1 | 1 | 0 | 0 | (5,4,3) | G0/G1 ( → G2) | C8, Cdc20=0, CyclinB=0 |
| Cdc25c=0 |  | 1 | 1 | 0 | 0 | 0 | 0 | 0 | 1 | (4,7,4) | G2 | Cdk1=0 |
| Cdk1=0 |  | 1 | 1 | 0 | 1 | 0 | 0 | 0 | 1 | (3,8,5) | G2 | C6_1/P5, C7_1 |
| CyclinB=0 |  | 1 | 0 | 0 | 1 | 1 | 1 | 0 | 1 | (4**,**5**,**4) | G2 | C6_2, C7_2 |
| CyclinA=1 |  | 1 | 0 | 1 | 1 | 0 | 1 | 1 | 1 | (1,4,5) | (G2 → ) SAC | C12_1, Cdh1=0, Cdc25c=1 |
| pAPC=1 |  | 1 | 1 | 0 | 1 | 0 | 1 | 1 | 1 | (2,7,6) | G2 ( → SAC) | CyclinA=0 |
| Cdc20=0 |  | 0 | 1 | 0 | 1 | 0 | 1 | 1 | 1 | (2,5,8) | SAC | C11/P2’ |
| Cdh1=0 |  | 0 | 0 | 1 | 0 | 0 | 0 | 1 | 1 | (4,3,4) | SAC → G0/G1 | C14, UbcH10=1 |
| Cdc20=1 | Tri-stable | 0 | 0 | 1 | 0 | 1 | 0 | 1 | 1 | (5,2,3) | G0/G1 | C12_3, C13 |
| Cdc20=1 |  | 0 | 0 | 1 | 0 | 1 | 0 | 1 | 0 | (6,1,2) | G0/G1 | C13 |
| Cdc20=1 |  | 0 | 0 | 1 | 0 | 1 | 0 | 0 | 1 | (6,3,2) | G0/G1 | C12_3 |
| CyclinB=1 | Bistable | 0 | 1 | 1 | 1 | 0 | 1 | 1 | 1 | (1,4,7) | SAC | C10, C12_2 |
| CyclinB=1 |  | 0 | 1 | 0 | 0 | 1 | 0 | 0 | 1 | (6,5,4) | G0/G1 ( → G2) | C12_2 |

**Supplementary Table S5.** **Phase Switch Oscillator states closest to Phase Switch attractors**

*Top:* The three fixed point attractors of the Phase Switch. *Middle:* the states of the Phase Switch Oscillator (wherein the states Wee1=Mad2=0 and Cdc25A=1, highlighted in gray, are fixed) that most closely approach the Phase Switch attractors under asynchronous update. The states closest to the G2 and SAC attractors are visited by part of the trajectories of the complex attractor (see Figure 4 and Figure 5). *Bottom:* the states that almost every asynchronous trajectory of the complex attractor will cross and are as close as possible to one of the three Phase Switch attractors. These three states also lie along the synchronous limit cycle. The highlighted cells represent states that differ from the respective Phase Switch attractor. A shared feature of both most likely close-to-attractor states is that they have all the nodes that form the Cyc meta-node in the ON state, while the G2 Phase Switch attractor does not have Cdk1 turned on yet and the SAC attractor already has CyclinA turned off.

The rightmost column, labelled passing probability, represents the likelihood of complex attractor trajectories passing through the state given in the row. These are estimates based on the filtered complex attractor shown on Figure 4 for the middle column, and the backbone shown in Figure 5 for the bottom column. The actual probabilities are somewhat smaller due to small probability shortcuts, see Figure 5 and Supplementary Table S2. The G0/G1 state in the middle and bottom table is the same*.*

| The attractors of the Phase Switch | | | | | | | | | | | |  |
| --- | --- | --- | --- | --- | --- | --- | --- | --- | --- | --- | --- | --- |
|  | **Cdc25C** | **CyclinA** | **Cdk1** | **CyclinB** | **Cdh1** | **pAPC** | **Cdc20** | **UbcH10** | **Mad2** | **Wee1** | **Cdc25A** |  |
| **G0/G1** | 0 | 0 | 0 | 0 | 1 | 0 | 0 | 0 | 0 | 1 | 0 |  |
| **G2** | 1 | 1 | 0 | 1 | 0 | 0 | 0 | 1 | 0 | 1 | 1 |  |
| **SAC** | 1 | 0 | 1 | 1 | 0 | 1 | 0 | 1 | 1 | 0 | 0 |  |
| The states of the Phase Switch Oscillator closest to the attractors of the Phase Switch | | | | | | | | | | | | Passing probability |
| **G0/G1** | 0 | 0 | 0 | 0 | 1 | 0 | 0 | 0 | 0 | 0 | 1 | 1.0 |
| **G2** | 1 | 1 | 0 | 1 | 0 | 0 | 0 | 1 | 0 | 0 | 1 | 0.33 |
| **SAC** | 1 | 0 | 1 | 1 | 0 | 1 | 0 | 1 | 0 | 0 | 1 | 0.55 |
| The most likely closest states of the Phase Switch Oscillator | | | | | | | | | | | |  |
| **G0/G1**  **(8,3,2)** | 0 | 0 | 0 | 0 | 1 | 0 | 0 | 0 | 0 | 0 | 1 | 1.0 |
| **Post-G2 (2,7,6)** | 1 | 1 | 1 | 1 | 0 | 0 | 0 | 1 | 0 | 0 | 1 | 0.89 |
| **near-SAC (1,6,7)** | 1 | 1 | 1 | 1 | 0 | 1 | 0 | 1 | 0 | 0 | 1 | 0.94 |

**Supporting References:**

S1: Perry, J. A., & Kornbluth, S. (2007). Cdc25 and Wee1: analogous opposites? *Cell division*, *2*(1), 12.
