## Supplemental Figures for "A feedback loop of conditionally stable circuits drives the cell cycle from checkpoint to checkpoint"

**Supplementary Figure S1. The limit cycle of the Phase Switch Oscillator under synchronous update**. Each column of squares indicates the states of the node written below the column. Each row corresponds to a state of the system. To use an unambiguous identifier that is more economical than indicating the state of all 8 noes, we describe the state by its overlap with the three attractors, in the order (G0/G1, G2, SAC). Each pair of successive rows (from top down) indicates a single synchronous update, i.e. applying the regulatory functions on the first state gives the second state. A dark grey square indicates the ON (1) state of the node indicated below the column and white means OFF (0).

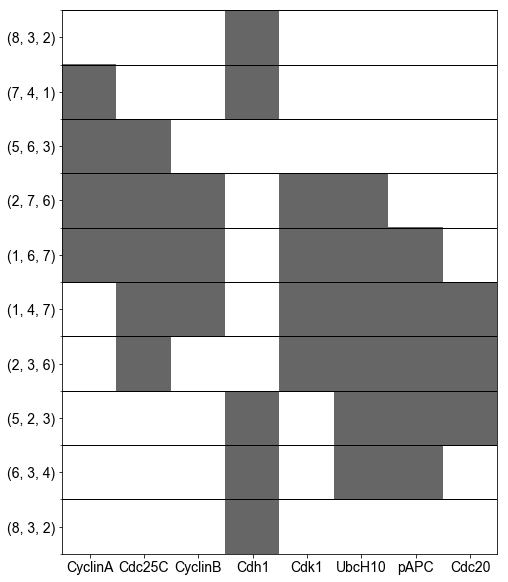

**Supplementary Figure S2. The expanded network of the Phase Switch.** The virtual nodes whose state is fixed in the Phase Switch Oscillator are shown in green.

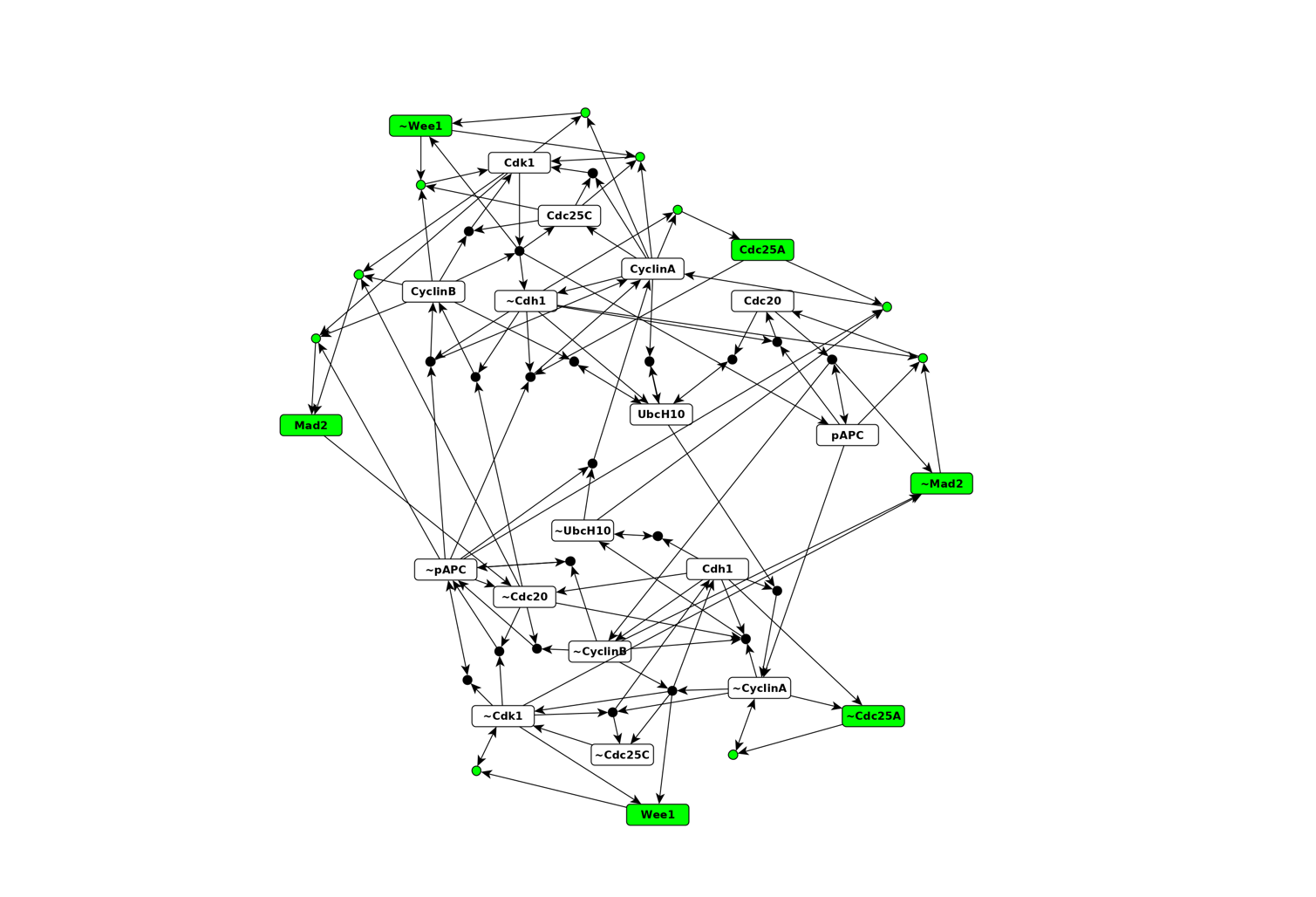

**Supplementary Figure S3. The distribution of the duration of the sustained on or off state of each node on the complex attractor of the Phase Switch Oscillator**. The distributions were obtained from an extensive number of trajectories of the system on the complex attractor, where each node alternates between being on and off. Each row corresponds to a node. The first figure in the row indicates the distribution of how long this node is on (with the median in red), the second figure indicates the duration of how long this node is off, and the third figure indicates the distribution of the duration of a consecutive on and off period. If the complex attractor were a deterministic cycle, the duration of a consecutive on and off period for any node were 16 time steps (i.e. each of the 8 nodes turning on once and turning off once). The observed medians are very close to 16. The split between the on and off periods is even (median on and off duration of 8) for five nodes and more asymmetric for three nodes, namely Cdc20, CyclinB, UbcH10. These three nodes also exhibit a slightly asymmetric pattern in the synchronous limit cycle, namely a 3 to 6 split of the nine steps as compared to the 4 to 5 split of the rest of the nodes (Figure 4).

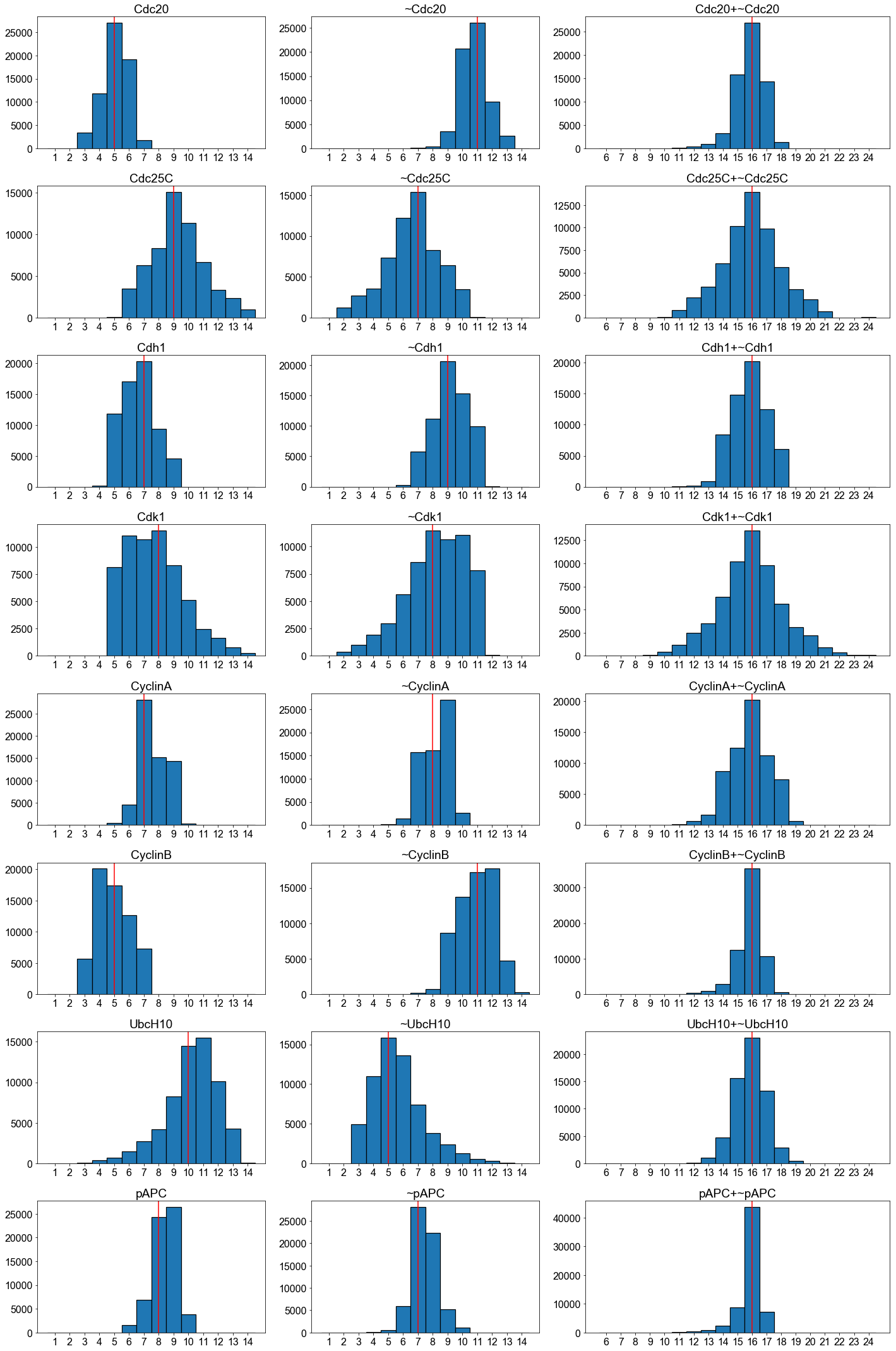

**Supplementary Figure S4. The expanded network of the Phase Switch Oscillator embodies the logic relationships that drive its oscillating behavior.** Colored nodes and edges highlight two characteristic subgraphs of the expanded network. The green nodes and edges indicate the subgraphs corresponding to the positive feedback loop between Cdk1 and Cdc25C (bidirectional edge in Figure 3 bottom panel). There are two overlapping cycles (i.e. closed paths with non-repeating virtual or composite nodes), both of length four. Each cycle involves Cdk1, Cdc25C and two composite nodes (one shared by both cycles). These cycles indicate that the positive feedback can only sustain the on (1) state of Cdk1 and Cdc25C if CyclinB (for one of the cycles) or both CyclinA and CyclinB (for the other cycle) are also simultaneously on. There is a consistent cycle formed by ~Cdk1, ~Cdc25C and a composite node that receives input from ~CyclinA; this means that Cdk1 and Cdc25 can simultaneously sustain their off (0) state if CyclinA is also off. The orange nodes and edges highlight the subgraph that corresponds to the negative feedback loop (bidirectional edge) between CyclinA and UbcH10 (Figure 3 bottom panel). This subgraph is a cycle of length eight; it contains both virtual nodes of CyclinA and UbcH10 as well as four composite nodes. In general, negative feedback loops result in a cycle in the expanded network involving both states of the involved nodes; we call this type of cycle inconsistent cycle. Positive feedbacks form two disjoint (groups of) cycles. Each of these cycles is consistent. The disjoint cycles have opposite states and can have different conditions.

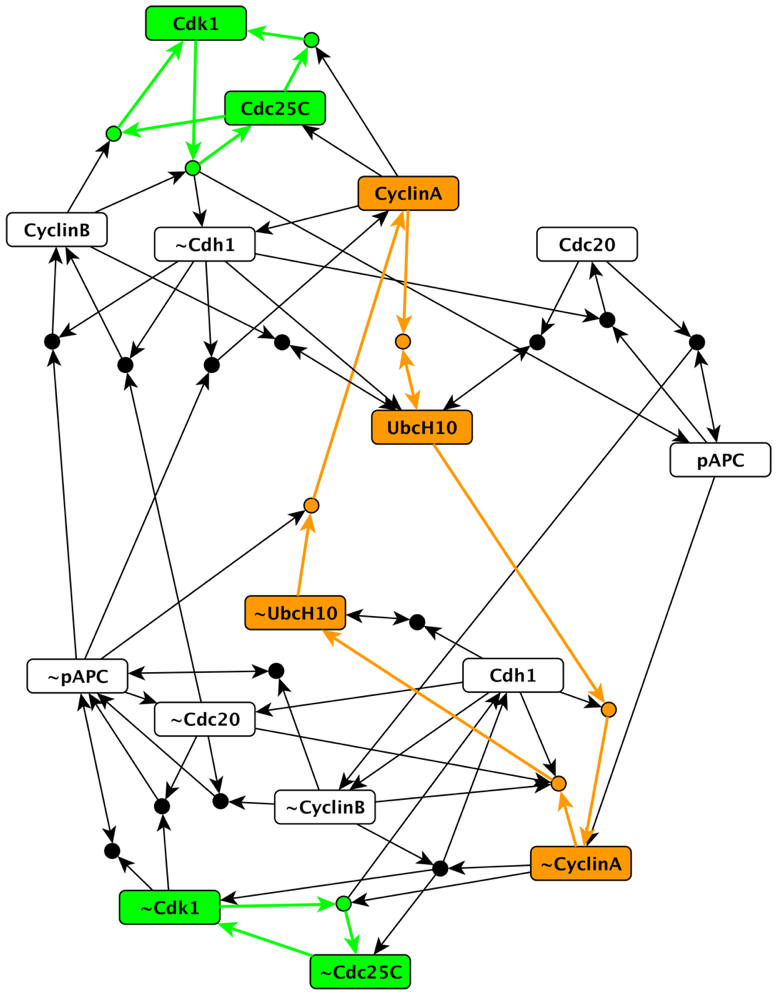

**Supplementary Figure S5. Distribution of the length of consistent (bottom) and inconsistent (top) cycles on the expanded network**. As illustrated by the green nodes and edges in Supplementary Figure S4, consistent cycles correspond to positive feedback loops in the regulatory network. They are the building blocks of conditionally stable motifs. We define inconsistency as the involvement of two opposite states of the same node, either as virtual nodes or as determinants of a composite node. Inconsistent cycles are akin to traversing a negative feedback loop of the regulatory network twice (as illustrated by the orange nodes and edges in Supplementary Figure S4).

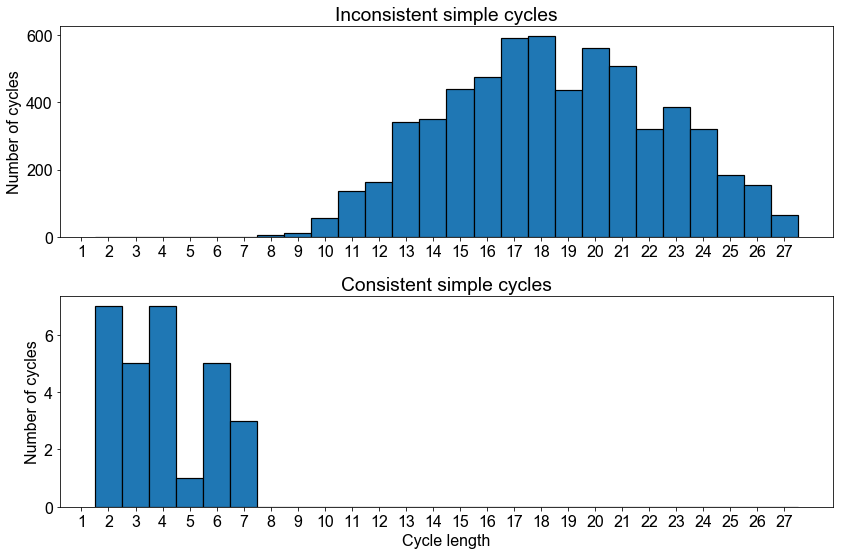

**Supplementary Figure S6. The virtual nodes of the expanded network contribute unequally to the oscillating motif.**  The weights of the edges among virtual nodes indicate the inverse of the probability of sufficiency (see Methods). The edges with high sufficiency (weight below 2) correspond to pairs of virtual nodes that are connected directly or through multiple composite nodes.  Virtual nodes marked by the node names preceded with ~ indicate the OFF state of the corresponding node. Node color represents the betweenness centrality, which is also indicated numerically beside each node. A darker shade indicates a higher betweenness centrality.

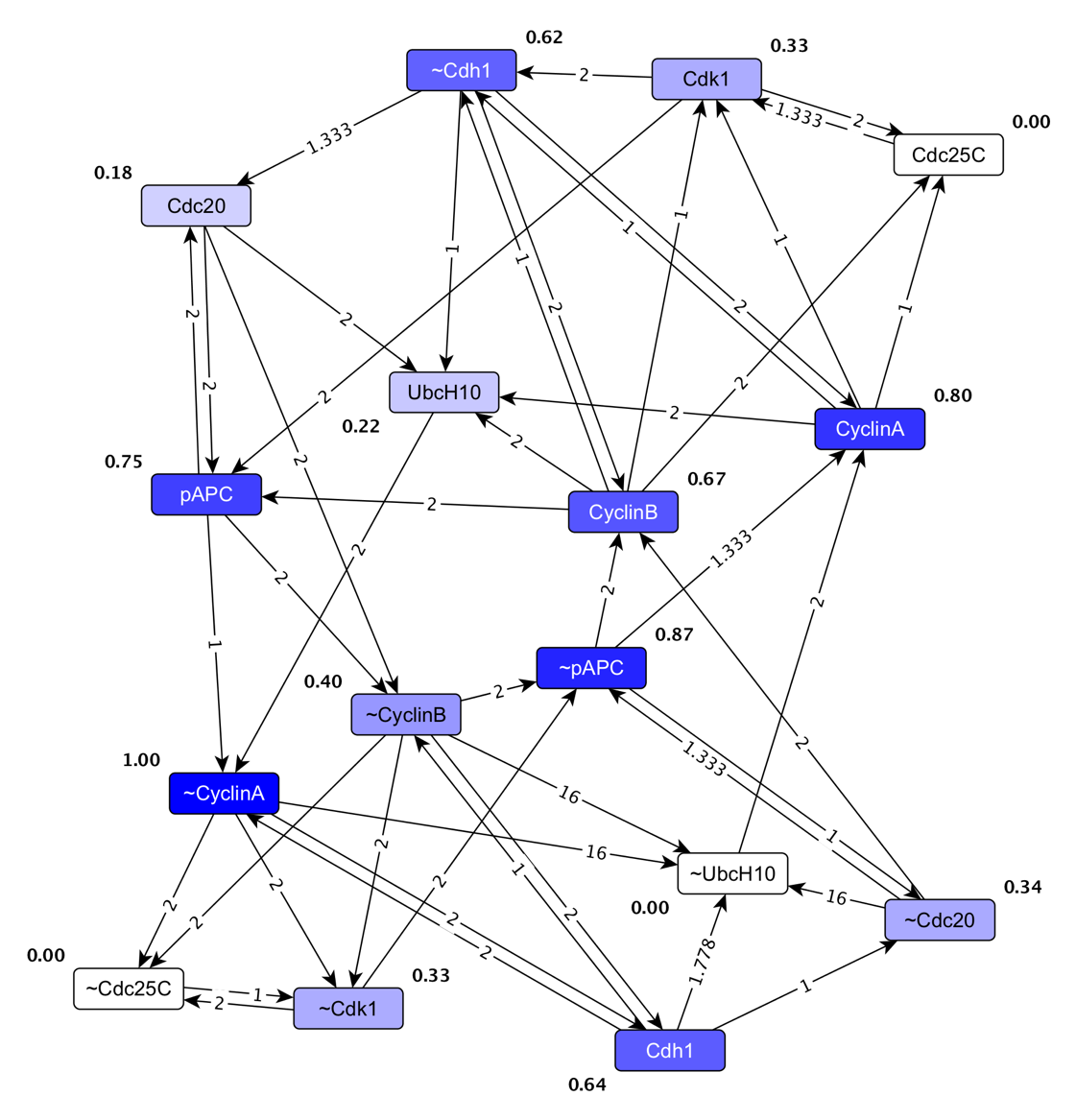

**Supplementary Figure S7. Illustration of a conditionally stable motif causing its own destabilization**. The conditionally stable motif C0, highlighted in green, is conditioned on ~CyclinA, emphasized with the dashed outline. The yellow nodes are the logical domain of influence (LDOI) of C0, i.e. states that will stabilize as long as C0 is stable. The on state of CyclinA, highlighted with orange, is also part of the LDOI but marks the destabilization of C0.

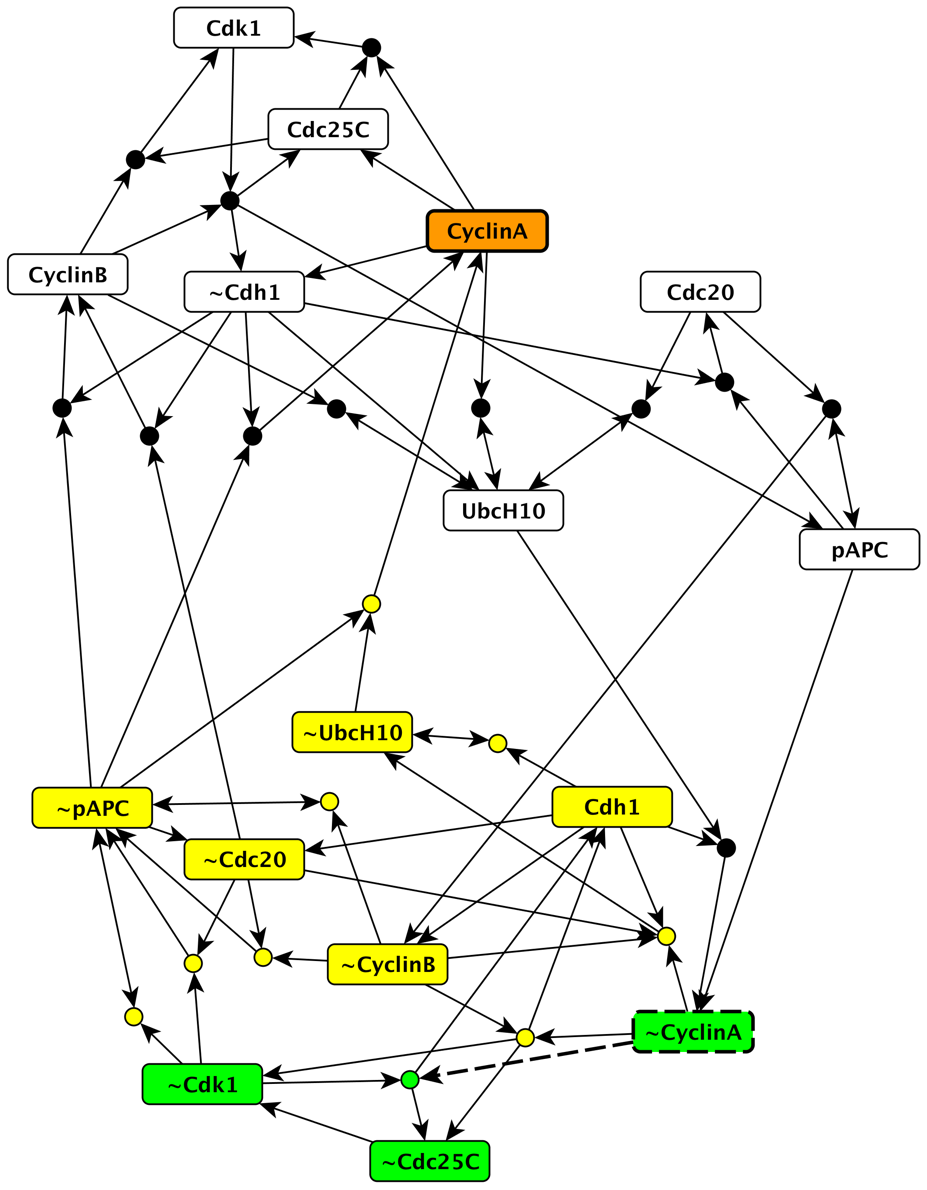

**Supplementary Figure S8. A sequence of sustained states of CyclinB can induce a sequence of transitions between attractors that mimics the cell cycle progression.**

The results are the average of 1000 simulations using general asynchronous update with random initial conditions. The state of the system is characterized by the overlap with the three attractors of the Phase Switch, where the maximal overlap is seven (the value of CyclinB is not included in calculating the overlap). In this sequence, CyclinB is manipulated to drive the system ensemble from random initial states through two cycles of the phase sequence G2 → SAC → G0/G1. A grey background indicates that CyclinB is forced off, while a white background indicates that it is forced on. Locking CyclinB off yields the G2 attractor of the system, and when CyclinB is subsequently forced on (t=150), the system converges into the SAC state. At the next time CyclinB is kept off (t=300), the system transiently overlaps with the G0/G1 attractor (indicated by the blue peak), but eventually converges to the G2 state. Keeping CyclinB on again (t=450) yields the SAC state. If CyclinB is subsequently off for a short duration only (t=600 to t=610), the system can converge to the G0/G1 attractor in the majority of trajectories. This is because the SAC-like state has CyclinA = 0, which, together with CyclinB = 0, is sufficient for Cdh1 = 1, Cdc25c = 0, and Cdk1 = 0. Furthermore, Cdh1 = 1 is sufficient for Cdc20 = 0, which is itself sufficient for pAPC = 0, making all nodes turn off.

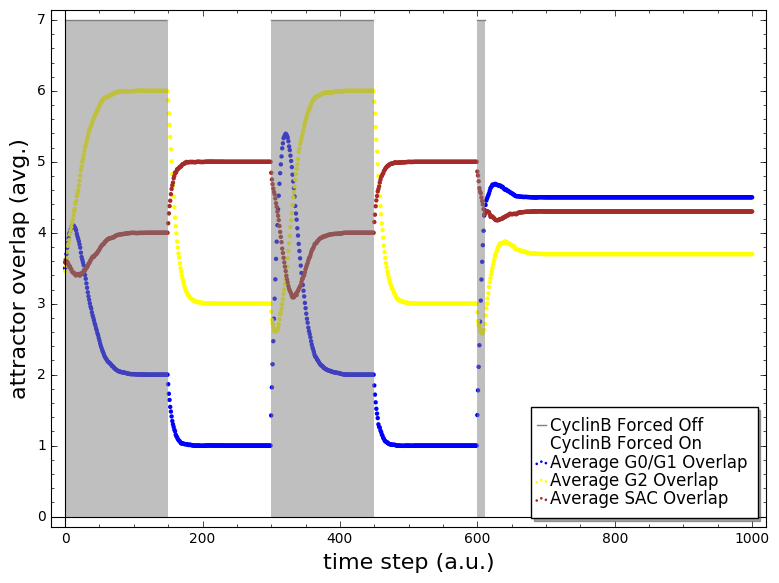

**Supplementary Figure S9. The cycle graph constructed for the Phase Switch Oscillator**. Each node of this graph represents a positive feedback loop of the regulatory network with node states (top row of each node label) that become self-sustaining when the state of certain other nodes (bottom row of each node label) is held fixed. As in previous figures the OFF state of a node is represented by a ~ preceding the respective node name. An undirected edge between nodes of the cycle graph indicates that the nodes states of the two cycles and their associated conditions are mutually compatible and non-disjoint. Every node and every consistent connected subgraph of the cycle graph corresponds to a conditionally stable motif. Note that in this case, all connected subgraphs are consistent. The largest connected subgraphs correspond to the meta-nodes Cyc (a subgraph of 10 cycles in the top left), ~Cyc (an 8-cycle subgraph in the top right), ~pAPC/Cdc20 (a four-clique in the bottom right), UbcH10 (triangle in the bottom left), pAPC/Cdc20 (two cycles connected by an edge) and ~UbcH10 (single cycle).

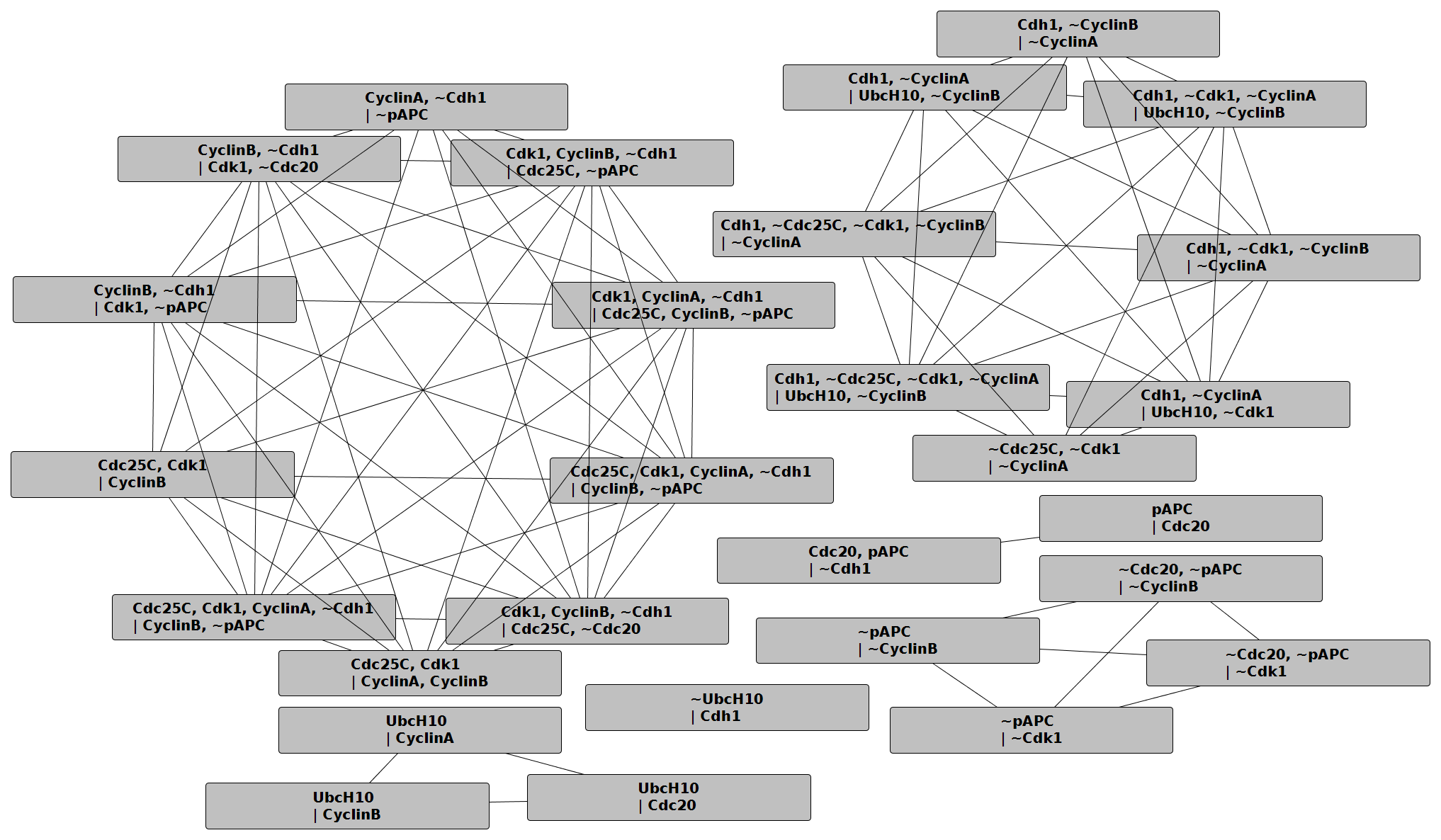
